# Pervasive cryptic selection in the human noncoding genome

**DOI:** 10.64898/2026.06.09.731256

**Authors:** Swetha Ramesh, Chenlu Di, Kirk E. Lohmueller

## Abstract

The prevailing dogma in evolutionary genetics holds that mutations within sequences that are conserved across a phylogeny are deleterious in those species, and mutations outside are neutrally evolving. Indeed, such comparative genomic approaches have estimated that mutations in approximately 5% of the human genome experience negative selection. However, sites that have biological function in certain lineages but not in others, i.e. functional turnover, may violate this assumption since these sites may be invisible to comparative genomic approaches. Thus, the extent of such cryptic, or hidden, negative selection remains elusive. Here, we developed a statistical test to detect cryptic selection in human polymorphism data. Applying our approach to simulated data shows that cryptic selection shapes the site frequency spectrum (SFS) and the statistical detection power depends on the proportion of mutations experiencing cryptic selection, the amount of sequence tested, and the sample size. We applied our method to polymorphism data from the 1000 Genomes Project, comparing variants in putatively functional noncoding regions to those in putatively neutral regions. We detected pervasive signals of cryptic selection in putatively functional regions, even after filtering out the top 70% of conserved sites. Using simulations with varying levels of cryptic selection, we estimated the extent of genome-wide constraint in the human genome. Our approximation suggests that mutations in at least 7% of the human genome are under negative selection, which is greater than the estimates from conservation-based methods, and that many of these mutations have escaped detection by comparative genomic methods. In sum, our results highlight the evolutionary dynamic nature of the noncoding genome and suggest the need to account for functional turnover when identifying putatively neutral variants for evolutionary analyses.

## INTRODUCTION

Early studies of molecular evolution focused on the protein-coding genome, which was the most exposed to negative selection (Reanney 1976). These studies found high levels of concordance in protein sequences across disparate species and reasoned that proteins were the main drivers of evolution. In 1969, Britten and Davidson challenged this paradigm by presenting an early model of gene regulation proposing that modifications of gene regulatory networks in higher organisms can be primary drivers of adaptation. Accordingly, in 1975, King and Wilson observed that human and chimpanzee protein sequences were more than 99% identical despite their significant anatomical and behavioral differences. They deduced that regulatory mutations outside of protein-coding genes play a more influential role in major evolutionary shifts than previously appreciated.

The advent of whole-genome sequencing technology enabled the creation of the first mammalian reference genomes, allowing for cross-species comparisons, i.e. comparative genomics (Mouse Genome Sequencing Consortium 2002; Miller et al. 2004). Higher sequence concordance between mammalian genomes made comparative genomics a feasible option for finding constrained noncoding sites (Miller et al. 2004). Constrained sites have high levels of similarity, i.e. are conserved, across organisms are indicative of negative natural selection having removed variation. Thus, such sites may have a biological function that relates to fitness (Miller et al. 2004). Candidate functional sites have been identified across a wide range of organisms, including mice, rats, dogs, and primates, and most of these sites lie outside of known coding regions and can be even more conserved than protein-coding genes (Shabalina 2001; Mouse Genome Sequencing Consortium 2002; Chiaromonte et al. 2003; Boffelli et al. 2003; Kirkness et al. 2003; Dermitzakis et al. 2003; Cooper et al. 2004; Bejerano et al. 2004; Miller et al. 2004; Cooper et al. 2005; Siepel et al. 2005; Birney et al. 2007; Asthana, Roytberg, et al. 2007; Parker et al. 2009; Garber et al. 2009; Pollard et al. 2010; Lindblad-Toh et al. 2011; Christmas et al. 2023). Many statistical approaches have also been developed to identify these regions with greater accuracy and precision (Margulies et al. 2003; Cooper et al. 2005; Siepel et al. 2005; Asthana, Roytberg, et al. 2007; Pollard et al. 2010; Arneson and Ernst 2019). These studies have revealed that around 3-8% of the human genome is under constraint, with some estimates being as high as 10-12% (Shabalina 2001; Asthana, Roytberg, et al. 2007; Parker et al. 2009; Garber et al. 2009; Christmas et al. 2023).

Despite the power of comparative studies in identifying sites with potentially deleterious mutations, such studies are unable to directly characterize the biological function at these sites. The ENCODE project, launched in 2003, has been the largest undertaking to address this issue by validating and classifying functional sites via biochemical assays. These assays revealed a surprisingly high proportion of the genome affects biological function, implicating 80.4% of the genome as functionally active (The ENCODE Project Consortium 2012). This estimate has been controversial, as it appears to be incompatible with the estimates predicted by comparative genomics studies. While criticisms have pointed to overly broad definitions of functionality (Doolittle 2013; Graur et al. 2013), another possible explanation for this discordance could be that portions of the genome may be biologically functional but not necessarily under evolutionary constraint (Doolittle 2013; Graur et al. 2013; Kellis et al. 2014). A related, and perhaps more nuanced explanation, is that noncoding sequences could also appear to be unconstrained due to functional turnover (Ponting and Hardison 2011). Sites under functional turnover have biological function in certain lineages but not in others due to changes in the regulatory networks and/or noncoding sequences over evolutionary time. In a scenario with functional turnover, mutations at noncoding sites may be under negative selection in certain lineages when they are biologically functional but are neutrally evolving in other lineages where they are not biologically functional. Thus, sites undergoing functional turnover may be invisible to comparative genomic approaches, given that data from species where the site is under negative selection and species where it is neutrally evolving are aggregated together. We refer to selection that is undetectable by comparative genomic approaches as cryptic selection.

Functional genomic studies have identified instances in support of functional turnover in mammalian genomes. For example, chromatin immunoprecipitation experiments revealed that in the liver, humans and mice differ in 41-89% of transcription factor binding locations, and less than 1% of enhancers are conserved across ten placental mammals (Young 2016). Several population genetics studies have also offered various lines of evidence for functional turnover over evolutionary time. Dermitzakis and Clark (2002) showed that transcription binding sites functionally conserved in humans and mice showed high sequence divergence, and they predicted that 32-40% of functional binding sites in humans are nonfunctional in mice. King et al. (2005) tested the predictive ability of two conservation-based metrics on a set of cis-regulatory modules (CRMs) in the human HBB locus. The two metrics were able to correctly identify only 50-60% of the CRMs, although 19 out of 23 tested CRMs were present in the multiple-species alignment used. Meader et al. (2010) and Rands et al. (2014) developed a method that used the distribution of distances between indels in the human genome to find regions under constraint. They observed an inverse relationship between the amount of pairwise sequence divergence and the amount of inferred mammalian constrained sequence. Ward and Kellis (2012) studied genome-wide constraint in humans for different partitions of ENCODE regulatory regions. They found non-conserved but functionally active ENCODE regions exhibited lower levels of genetic variation, implying higher levels of negative selection than functionally inactive ENCODE regions. Huber et al. (2020) used simulations to show that the GERP score, a comparative genomic constraint score, has little power to predict sites under lineage-specific negative selection when aggregating sequences from a wide range of species. They fitted models with and without functional turnover to the empirical GERP score distribution for humans and found that the best fitting model incorporated turnover. Christmas et al. (2023) estimated a lower bound of 10.2% of sequence under constraint using an alignment of 240 placental mammals–one of the largest studies of this kind to date. They noted that including more closely related species in the alignment correlates with increased estimates of genome-wide constraint. They also note that simple utilization of PhyloP scores (another constraint score) are underpowered at identifying weakly constrained and lineage-specific constrained sites. Di et al. (2025) explored the distribution of fitness effects (DFE) of noncoding mutations in the human genome. They found that 50% of mutations within the top 5% of sites most conserved in primates, but not mammals, are inferred to be under negative selection. These studies all suggest that sites can undergo changes in selective effects over evolutionary time, but comparative genomics methods are underpowered to capture this phenomenon.

When considering functional turnover, estimates of genome-wide constraint increase to 8-15% (Smith et al. 2004; Rands et al. 2014; Di et al. 2025) relative to estimates from comparative genomic approaches. However, the very nature of functional turnover limits its identification and obfuscates the true amount of sequence undergoing negative selection. Despite this, non-conserved noncoding regions are routinely assumed to be neutrally evolving in population genetic studies, such as those inferring demography or the strength of background selection. Residual, uncaptured sites undergoing negative selection can be misinterpreted as a signature of recent population growth in demographic inference (Gazave et al. 2013) or can lead to an overestimate of background selection (Charlesworth and Jensen 2021). However, the optimal filters for removing sites where mutations are directly under selection are not always clear. Consequently, different studies have taken varied approaches to define a set of neutral sites. For example, the popular Out-of-Africa model from Gravel et al. (2011) was inferred using ∂a∂i, a maximum likelihood-based method for demographic inference using diffusion approximations (Gutenkunst et al. 2009). They used sites within noncoding DNA sequence nearby various autosomal genes without excluding any known conserved sites. Excoffier et al. (2013) introduced fastsimcoal2 and applied their method to infer demography for African, European, and African-American populations using intergenic SNPs outside CpG islands but did not exclude other types of conserved sequence. Phung et al. (2016) investigated the extent to which background selection in the ancestral population can affect levels of neutral divergence in a wide range of taxa. They identified putatively neutral sites using stringent filtering criteria, including removing sites within the top 10% of GERP scores, and observed a negative correlation between neutral divergence and the functional content overlapping neutral sites. However, when filtering was tightened to exclude the top 15% of conserved sites, the previously significant correlation in humans and rodents disappeared. Buffalo and Kern (2024) created a theoretical model to quantify background selection in the human genome. They tested their model using simulations and applied it to human genome data by estimating levels of diversity, substitution rates, and the DFE. For their model, they defined putatively neutral sites as sites at least 200bp outside of exonic and coding regions. However, they noted that the pairwise diversity of their neutral sites was much lower compared to sites within the bottom 70% of CADD scores, another conservation-based metric. In fact, they found that their predictive model underestimates pairwise diversity and substitution rates–a possible consequence of their neutral site identification criteria. These studies use sequence conservation as a proxy for negative selection, yet it is still unclear how conservation-based filtering approaches perform at removing variants under selection. If the majority of the noncoding, deleterious mutations occur in conserved regions, then comparative genomic filters should be sufficient. However, if a non-negligible number of deleterious mutations appear outside of conserved regions, more stringent filtering criteria may be needed.

Overall, the extent to which comparative genomic approaches detect most of the noncoding, deleterious mutations of the human genome remains unclear. Patterns of polymorphism within humans provide an alternate way to detect selection in the noncoding regions of the human genome. Numerous studies have suggested that mutations in conserved regions experience selection in humans (Asthana, Noble, et al. 2007; Asthana, Roytberg, et al. 2007; Bejerano et al. 2004; Birney et al. 2007; Boffelli et al. 2003; Chiaromonte et al. 2003; Christmas et al. 2023; Cooper et al. 2004; 2005; Dermitzakis et al. 2003; Di et al. 2025; Garber et al. 2009; Kirkness et al. 2003; Lindblad-Toh et al. 2011; Miller et al. 2004; Mouse Genome Sequencing Consortium 2002; Parker et al. 2009; Pollard et al. 2010; Ponting and Hardison 2011; Shabalina 2001; Siepel et al. 2005; The ENCODE Project Consortium 2012; The ENCODE Project Consortium et al. 2020). However, the complementary question–whether non-conserved regions are neutrally evolving–has not been fully addressed using polymorphism data.

Here, we test whether we can detect differences in patterns of polymorphism data in humans as summarized by the site frequency spectrum (SFS) when cryptic selection is present in non-conserved noncoding regions. The SFS is the distribution of allele frequencies in a population of interest, and it is a commonly used summary statistic in population genetics. We developed a likelihood ratio test (LRT) for homogeneity to compare SFSs generated under different conditions. We applied this test to forward-in-time simulations with 0%, 1%, 3%, and 5% of mutations in non-conserved noncoding sequence drawn from a DFE estimated from human conserved noncoding (CNC) sequence (Torgerson et al. 2009), emulating cryptic selection potentially present in the human genome. We find that SFSs for conditions with cryptic selection are significantly different from those without, and the magnitude of the difference varies depending on the number of individuals sampled and amount of sequence tested. We then apply our approach to human genetic variation in putatively functional and putatively neutral genomic regions, identifying strong signals of negative selection that are not captured by conservation scores. Using our simulations as benchmarks, we estimate that mutations in at least 7% of the human genome are under constraint. These results demonstrate that overlooked cryptic selection in the noncoding genome has a tangible effect on the SFS and may impact inferences in downstream applications.

## RESULTS

### Simulation setup

We first characterized the impact of cryptic selection on genetic variation in simulated data using SLiM, a forward-in-time simulation framework (Haller and Messer 2023). Each simulation replicate comprised a 20 Mb genome that included randomly placed coding segments, conserved noncoding (CNC) segments, and a large, contiguous neutral segment. Any sequence not within these three categories was considered background noncoding (BGNC), i.e. non-conserved noncoding, and encompassed a majority of the simulated genome.

BGNC sequence is often assumed to be neutrally evolving in many downstream population genetic inference approaches. We included negative selection in BGNC regions in our simulations and tested whether it can be detected in the SFS. BGNC regions were simulated under conditions of 0%, 1%, 3%, or 5% of mutations with selection coefficients drawn from a DFE estimated from CNC sequence in humans (Torgerson et al. 2009), and the rest of the mutations were neutral. We hereby refer to this DFE as the ncDFE. Mutations subject to the ncDFE were deleterious (|*s*|>10^−5^) with a probability of 26% (Supplementary Table 1). Deleterious mutations arising in BGNC sequence embody the cryptic selection that may be present in the human noncoding genome due to functional turnover. All mutations in the CNC regions were subject to the ncDFE. Coding regions were simulated to have a 2.31:1 ratio of nonsynonymous (deleterious) to synonymous (neutral) mutations, following Kim et al. (2017). All mutations arising within the large neutral segment were selectively neutral. Population size changes in all simulation replicates followed the European demographic model estimated by Kim et al. (2017).

### Cryptic selection alters the site frequency spectrum in simulations

We obtained SFSs for sample sizes of 10, 100, and 1000 haploid genomes from each of our simulation replicates. A qualitative comparison of the proportional and residual SFSs for the BGNC and CNC regions shows a clear skew towards low-frequency variants as the amount of sequence under cryptic selection increases (Figure 1). The skew in the BGNC sequence is pronounced regardless of sample size (Figure 1A, Supplementary Figure 1A). These trends are expected and arise from the direct effects of negative selection acting on the mutations in the BGNC regions. Notably, we also observe the skew in the SFSs for the CNC regions as the amount of cryptic selection in BGNC regions increases. This is due to the effects of background selection from linked deleterious mutations in BGNC (Figure 1B, Supplementary Figure 1B). The CDS and neutral regions also exhibit an excess of rare variants and a deficit of common variants, just like the CNC regions, but it is less obvious at smaller sample sizes (Supplementary Figure 1C-D).

**Figure 1.**
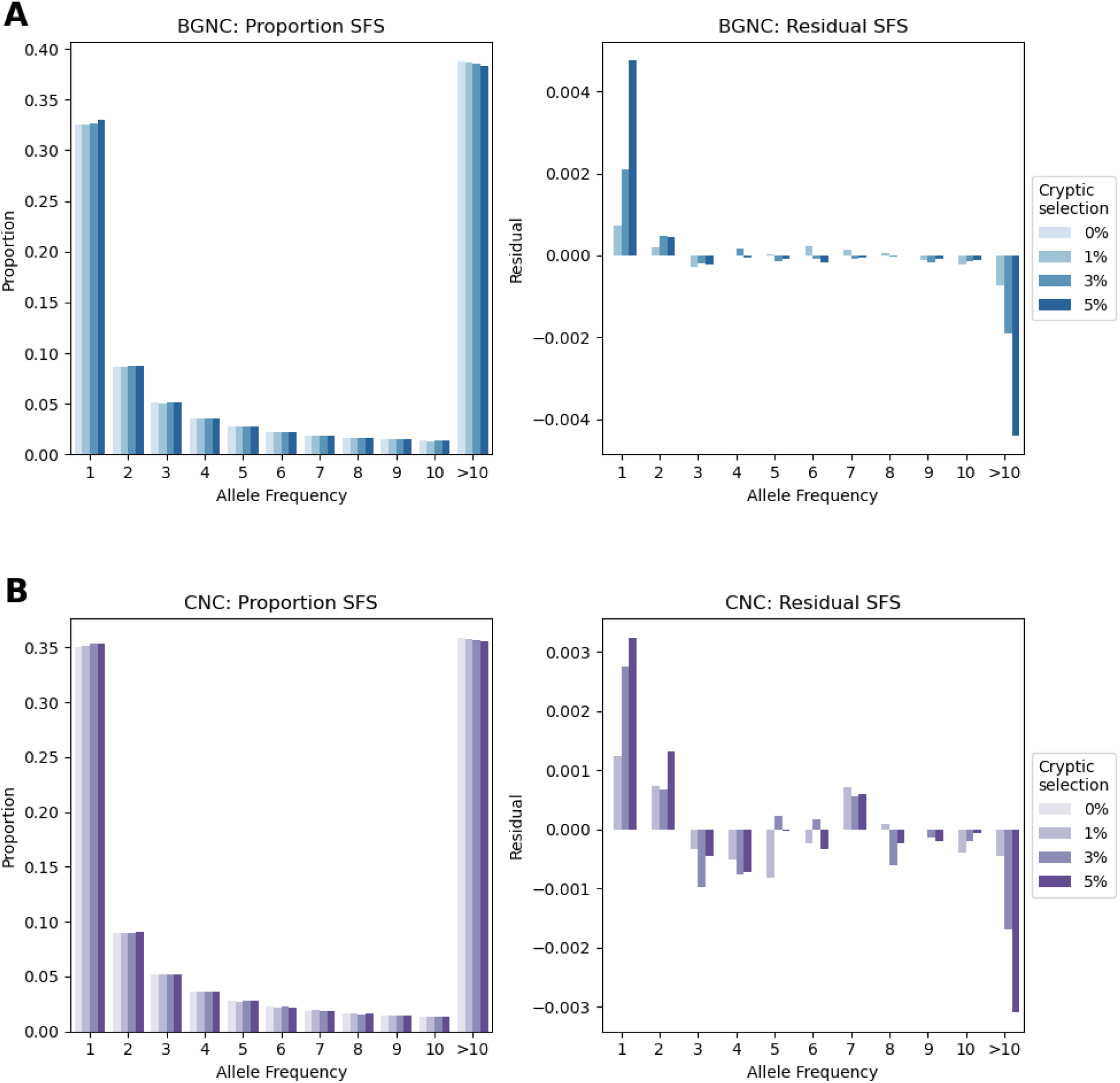
SFSs for 100 sampled genomes for BGNC and CNC regions from simple simulations with 0%, 1%, 3%, and 5% of sites under the ncDFE. On the left of each panel is the proportion SFS, which shows the proportion of variants in each allele frequency bin for each cryptic selection condition. On the right of each panel shows the residuals, calculated by subtracting the bin values for the 0% condition from the other conditions. SFSs for BGNC (A) and CNC (B) regions are shown here.

### LRT for homogeneity to compare SFSs

We developed an LRT for homogeneity to statistically detect the effects of cryptic selection in the SFS. Essentially, we compare one SFS that consists of variants that we are confident are neutrally evolving with another SFS that contains variants potentially under negative selection. We only considered segregating variants in this test. The number of SNPs in each bin of the SFS can be considered a Poisson random variable (Sawyer and Hartl 1992), thus we used a Poisson likelihood function. Under the null hypothesis, for a sample size of *n* haploid genomes, the two SFSs, *X* = {*x*_1_, *x*_2_, …, *x*_*n*−1_} and *Y* = {*y*_1_, *y*_2_, …, *y*_*n*−1_}, are from the same condition (i.e. neutrality). Therefore, the rate parameters *λ* of the Poisson distributions will be the average of the two observed SFSs. The log-likelihood under the null hypothesis is as follows:

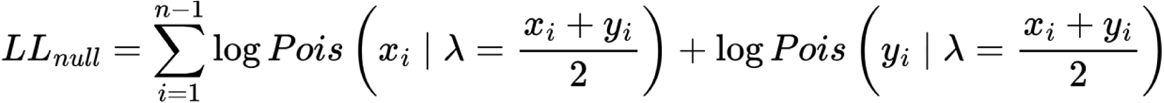

Under the alternative hypothesis, the two SFSs are from different conditions (i.e. neutrality and negative selection). Therefore, the rate parameters of the Poisson distributions for each SFS will be those from its maximum likelihood estimate, i.e. the SFS itself:

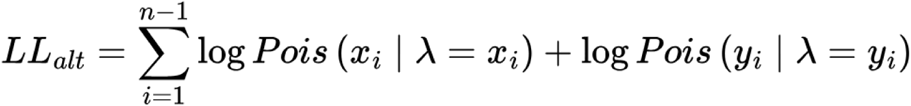

The LRT statistic is then calculated as follows:

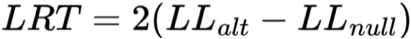

### Testing the LRT for homogeneity on simulated data

We quantitatively evaluated the performance of our LRT to detect cryptic selection by comparing the SFSs for BGNC regions from simulations with and without cryptic selection (Figure 2). More specifically, we compared SFSs of BGNC regions from simulations with 0% of mutations subject to the ncDFE to simulations with 1%, 3% and 5% of mutations subject to the ncDFE. To investigate the effects of cryptic selection across different sequence lengths (i.e. simulation subset sizes), we employed a bootstrap procedure to sample from our simulation replicates. We sampled a subset of SFSs from a set of simulations with cryptic selection and a set without. We summed the SFSs in each subset across SFS bins and compared the two aggregated SFSs using the LRT for homogeneity. This process was repeated until we obtained 1000 LRT statistics. We used the mean LRT statistic to determine statistical significance. We conducted this entire procedure for each strength of cryptic selection for multiple lengths of simulated sequence and sample sizes.

**Figure 2.**
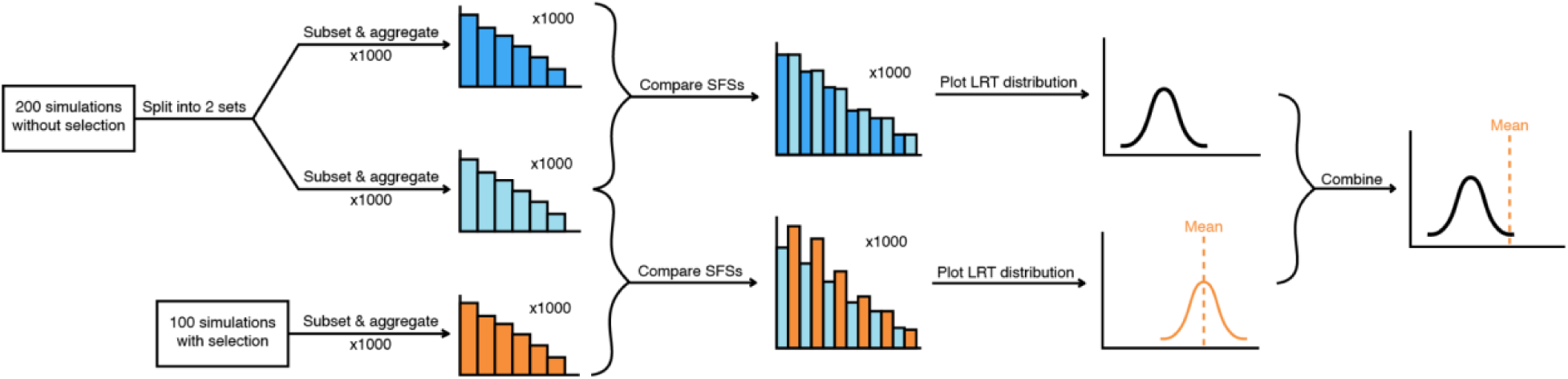
Flowchart of SFS comparison strategy. Sets of simulations are repeatedly sampled and aggregated. Aggregate SFSs for two sets of simulations without selection are compared using our LRT of homogeneity and the resulting LRT statistics comprise the null distribution. Aggregate SFSs for one set of simulations with selection and one without selection are compared in the same manner, and the resulting LRT statistics are averaged. The null distribution and mean LRT statistic are used together to determine whether there is a statistically significant difference.

We opted to create an empirical null distribution from our simulations without cryptic selection rather than rely on the asymptotic chi-square distribution of the LRT under the null hypothesis. Our rationale was that for all sequence lengths and sample sizes tested, there was a considerable rightward shift in the empirical null distribution of LRT statistics compared to the chi-square distribution (Supplementary Figure 2), likely because the assumptions of linkage equilibrium of the SNPs does not hold when considering large amounts of data (Bustamante et al. 2001; Zhu and Bustamante 2005). Similar to before, we generated two sets of simulations without cryptic selection and obtained their SFSs. We repeatedly sampled subsets of SFSs from each set of simulations, summed the SFSs for each subset, and then compared them using the LRT for homogeneity. The 1000 resulting LRT statistics comprised the empirical null distribution. We determined statistical significance by comparing this null distribution to the mean LRT statistics calculated from the comparison of simulations with and without cryptic selection.

The detection of cryptic selection varied based on its strength as well as the number of haploid samples and the length of simulated sequence tested (Figure 3). At 1 Gb and 600 Mb of simulated sequence (first and second rows of Figure 3), the mean LRT statistics for 3% and 5% of BGNC mutations under the ncDFE (equivalent to 0.782% and 1.30% of BGNC mutations under negative selection, respectively (Supplementary Table 1)) surpass the significance threshold of *α* = 0.05 across all three sample sizes. At 200 Mb (third row of Figure 3), the mean LRT statistics for only 5% of BGNC mutations under the ncDFE is significant at all three sample sizes. At 100 Mb (last row of Figure 3), the mean LRT statistics for 5% of BGNC mutations under the ncDFE is significant at only sample sizes of 10 and 100. The mean LRT statistics for 1% fails to reach significance for all the sample sizes and sequence lengths tested. This indicates we may not have enough data to have sufficient power to detect very weak cryptic selection. However, this level of cryptic selection is much lower than current estimates of constraint in the human genome, as 1% of BGNC mutations under the ncDFE is equivalent to 0.261% of BGNC sites under negative selection (Supplementary Table 1).

**Figure 3.**
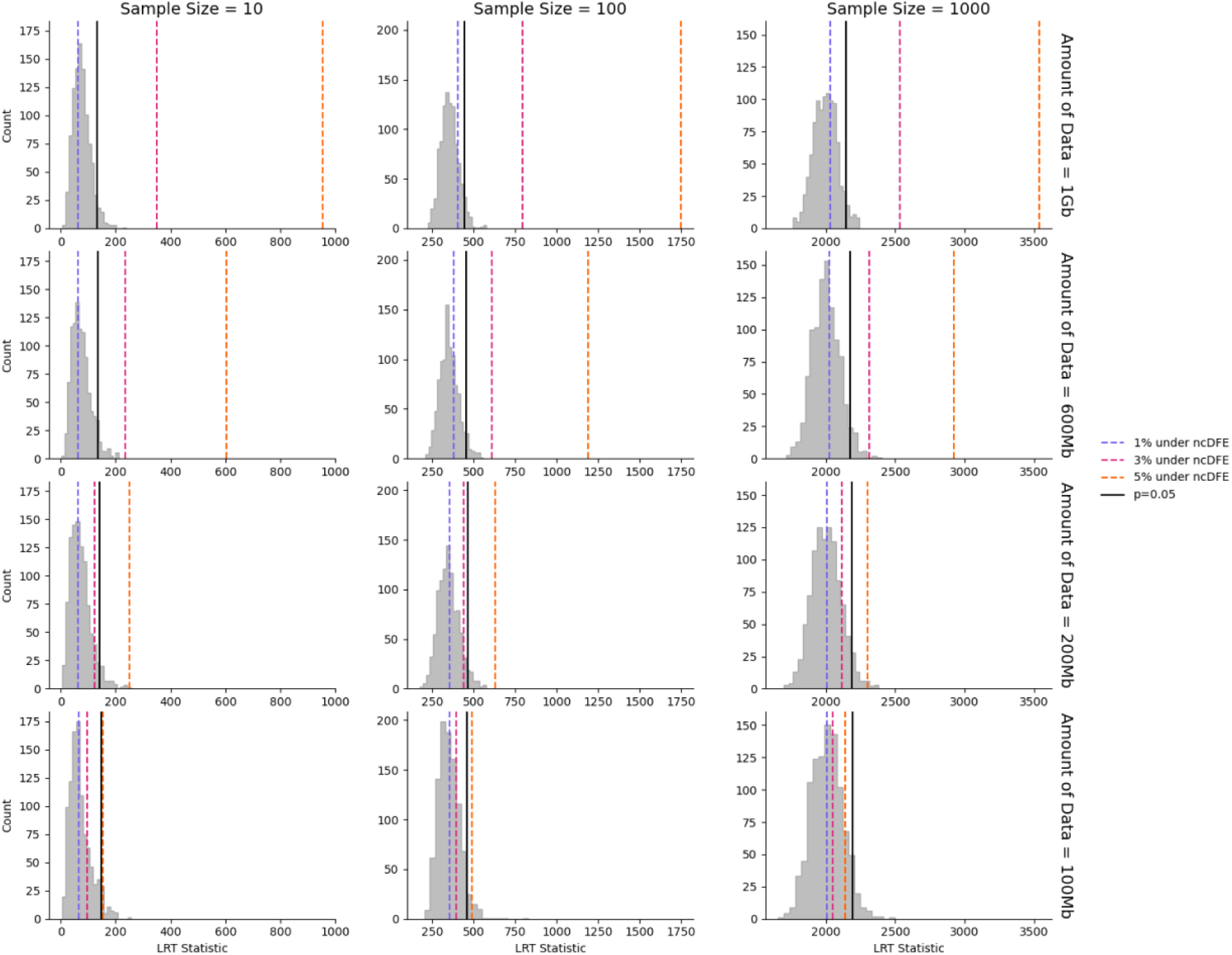
LRT plots for simple simulations with and without cryptic selection. Each row and column correspond to different total sequence lengths and sample sizes, respectively. Sample sizes are in terms of number of chromosomes. Within each plot, the gray histogram represents the empirical null distribution, i.e. the distribution of LRT statistics from comparing two sets of simulations with 0% cryptic selection. The black line marks the critical value of the empirical null distribution. The colored lines correspond to the mean LRT statistic from comparisons of simulations with no cryptic selection to simulations with 1%, 3%, or 5% of BGNC mutations subject to the ncDFE. Colored lines that lie past black lines denote statistically significant LRT comparisons.

Interestingly, we observed greater statistical power at a sample size of 100 haploid genomes, rather than at 1000 (Supplementary Figure 3). The theoretical intuition of Kallenberg et al. (1985) helps explain this non-monotonic behavior. They show that in likelihood ratio goodness-of-fit tests, increasing the number of bins has two opposing effects on power: it can increase the chi-squared distance between the null and alternative distributions, but can also increase the variance of the null distribution and thus the critical value. Although they assume a fixed total count and derive this logic for goodness-of-fit tests, which have known null probabilities, we can still extend their findings to our test for homogeneity. As the sample size, i.e. number of SFS bins, grows exponentially from 10 to 100 to 1000, the expected number of segregating sites grows only linearly (Watterson 1975). Furthermore, the signal of cryptic selection is concentrated in the low-frequency bins, as rare variants are enriched under negative selection (Figure 1). At a sample size of 10, there are too few bins to resolve the signal of cryptic selection, whereas a sample size of 1000 has too many bins, which inflates the variance of the null distribution, and thus the critical value. A sample size of 100 optimally balances both of these effects and yields the greatest observed power. Nonetheless, statistical power to detect cryptic selection generally increases with dataset size and the strength of cryptic selection, reflecting the increased ability to distinguish SFSs with and without cryptic selection.

The patterns seen in this set of simulations may, in part, reflect the simplified structures of the simulated genomes. Thus, we carried out additional simulations to determine whether these patterns persist in scenarios with complex, real-world genome structure, linkage disequilibrium (LD), and background selection. For each simulation replicate, we matched the locations of coding and CNC sites in the 20 Mb genome to those of human coding and CNC annotations within a 20 Mb region sampled from the human genome (see Methods). The rest of the sites in each simulated genome were classified as BGNC, which was simulated to undergo the same four proportions of mutations undergoing cryptic selection as in the previous simulations. All other aspects of our simulations remained the same as before.

We employed the same bootstrap procedure to compare simulations with and without cryptic selection for a range of sample sizes and simulated sequence lengths. As before, we created an empirical null distribution of LRT statistics by comparing two sets of simulations without cryptic selection. Then, we compared simulations with and without cryptic selection to obtain mean LRT statistics, which were used along with the empirical null distributions to investigate the ability of our LRT for homogeneity to detect cryptic selection (Figure 4). Here, empirical null distributions have larger means and variances compared to the previous simulations, likely due to the genome structures of the simulations better reflecting the variability seen in the human genome. At 1 Gb (first row of Figure 4), conditions where 3% and 5% of BGNC mutations under the ncDFE are significantly different from the null distribution at all sample sizes. At 600 Mb (second row of Figure 4), 5% continues to be significantly different at all sample sizes, and 3% falls past the significance threshold only for sample sizes of 10 and 100. This mirrors the pattern seen in the previous set of simulations, where the intermediate sample size of 100 tends to better capture the signal of cryptic selection than sample sizes of 10 and 1000 (Supplementary Figure 4). At 200 Mb (third row of Figure 4), the condition where 5% of the mutations in BGNC regions are from the ncDFE is significantly different at sample sizes of 10 and 100, but not 1000. At 100 Mb (fourth row of Figure 4), none of the levels of cryptic selection we tested were significantly different from the null distribution. Once again, 1% fails to surpass the significance threshold at any sample size and sequence length tested. Essentially, we observed the same patterns as the previous set of simulations, albeit slightly weaker. In these complex, realistic simulations, we detect differences in the SFSs with at least 200 Mb of sequence (Figure 4). Whereas in the previous, simpler simulations, we detect differences with at least 100 Mb of sequence (Figure 3). Nevertheless, the impact of cryptic selection on the SFS remains detectable by our LRT for homogeneity.

**Figure 4.**
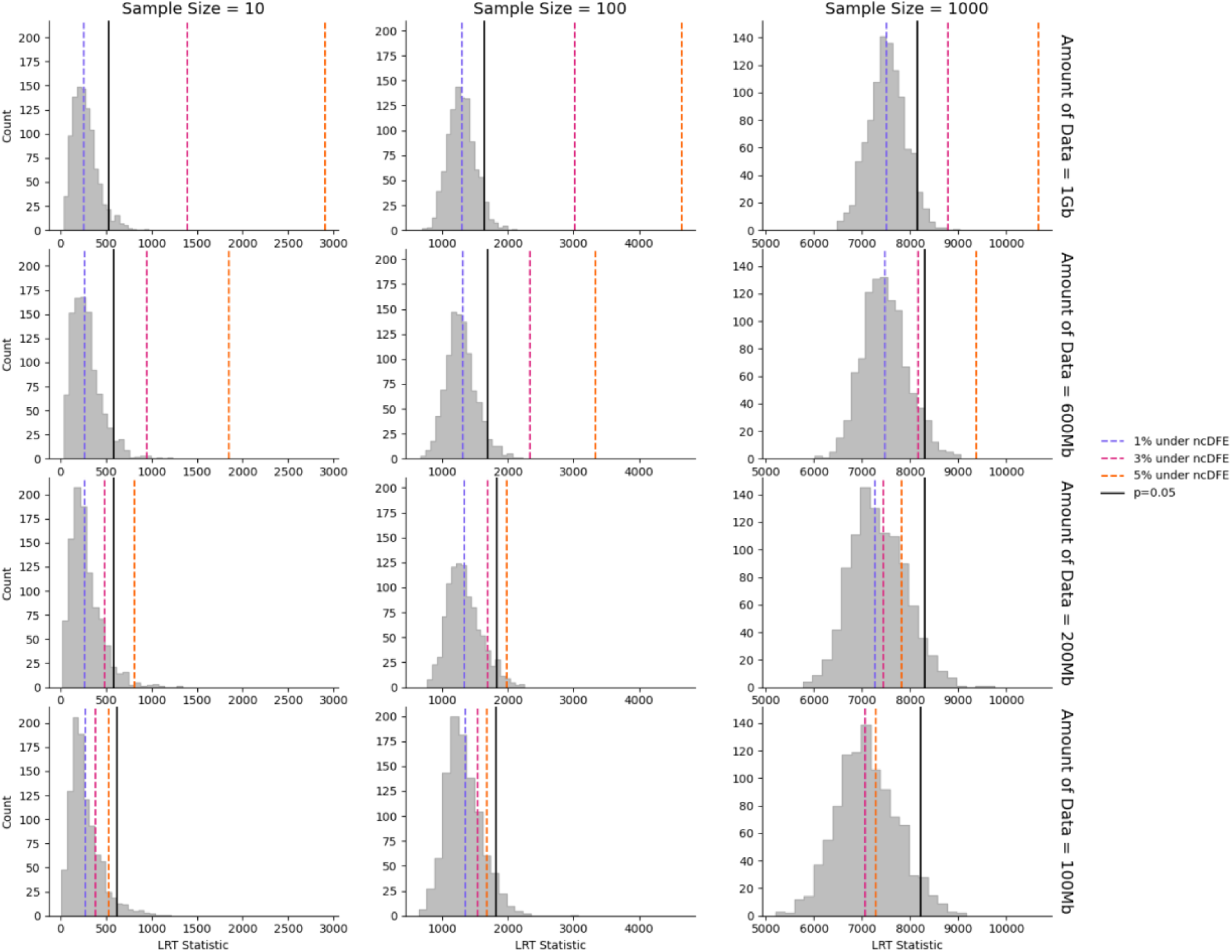
LRT plots for complex, realistic simulations with and without cryptic selection. Each row and column correspond to different total sequence lengths and sample sizes, respectively. Sample sizes are in terms of number of chromosomes. Within each plot, the gray histogram represents the empirical null distribution, i.e. the distribution of LRT statistics from comparing two sets of simulations with 0% cryptic selection. The black line marks the critical value of the empirical null distribution. The colored lines correspond to the mean LRT statistic from comparisons of simulations with no cryptic selection to simulations with 1%, 3%, or 5% of BGNC mutations subject to the ncDFE. Colored lines that lie past black lines denote statistically significant LRT comparisons.

Altogether, our simulations show that cryptic selection can have a considerable impact on the SFS. The efficacy of capturing its effects mainly depends on the sample size and amount of sequence tested. Genome structure can also influence its effectiveness, as evidenced by the weaker ability to detect cryptic selection in genomes of complex structure. Our LRT for homogeneity detects the effects of cryptic selection at a minimum of 0.781% of BGNC mutations under weak negative selection, equivalent to 3% of BGNC mutations under the ncDFE (Supplementary Table 1). Our results demonstrate that cryptic selection has the potential to significantly impact the SFS and, by extension, downstream inference approaches.

### Comparison of SNPs in human putatively functional and quiescent sequence reveal pervasive cryptic selection in the human genome

Next, we tested whether real human genomes show evidence of cryptic selection by applying our LRT for homogeneity to SNP data from the 1000 Genomes Project (1KGP). We first categorized sites as either “putatively functional” or “quiescent” using functional genomic annotations from ChromHMM (see Methods). Quiescent sequence has a paucity of transcriptional activity, histone modifications, and epigenetic marks and is thus considered functionally inactive (Hoffman et al. 2013). Therefore, it is more likely to be evolving neutrally. Putatively functional sequence is any noncoding sequence outside of quiescent sequence that may contain sites undergoing functional turnover and experiencing negative selection. Here, quiescent and putatively functional sequences serve as empirical analogs to simulations without and with cryptic selection, respectively. Thus, similar to before, we compared the SFSs from these two types of sequences to determine whether cryptic selection has shaped the SFS of putatively functional regions. We also tested how phylogenetic constraint can affect the detection of cryptic selection, as any negative selection occurring within CNC sequence may obscure SFS differences created by cryptic selection alone. We excluded sites, ranging from those within the top 10% to the top 70% of PhastCons scores, from both the quiescent and putatively functional datasets of sequences. See Supplementary Table 2 for the amounts of resulting sequence before and after excluding CNC sites.

Quiescent sequence is substantially less abundant in the human genome than putatively functional sequence (Supplementary Table 2). To ensure we were comparing equivalent amounts of sequence from each category across the different PhastCons filtering thresholds, we randomly downsampled quiescent and putatively functional sites to total 100 Mb each. We confirmed that this downsampling did not introduce any differential LD by inferring pairwise LD in each 100 Mb subset using PLINK (Supplementary Figure 5). While LD decay curves show a slight difference between putatively functional and quiescent sites (Supplementary Figure 5A), the cumulative distributions of r^2^ values for both datasets are nearly identical to each other (Supplementary Figure 5C), suggesting that downsampling did not introduce any substantial biases that may influence our downstream analyses.

We first investigated the SFSs from the putatively functional and quiescent datasets to determine how sample size and the amount of conserved sites excluded, i.e. PhastCons percentile thresholds, qualitatively affects patterns of genetic variation (Figure 5). The SFSs for putatively functional sites have a striking skew towards rare variants and away from common variants for sample sizes of 10 and 100 (first two columns of Figure 5) relative to the quiescent SFS. This skew persists even when excluding the top 70% of conserved sites (last row of Figure 5), a level at which presumably most deleterious sites would be excluded (Di et al. 2025). For a sample size of 1000 (third row of Figure 5), the skew away from common variants is clear but the skew towards rare variants is not as obvious. Broadly, putatively functional regions exhibit patterns consistent with negative selection, even when excluding evolutionarily conserved regions.

**Figure 5.**
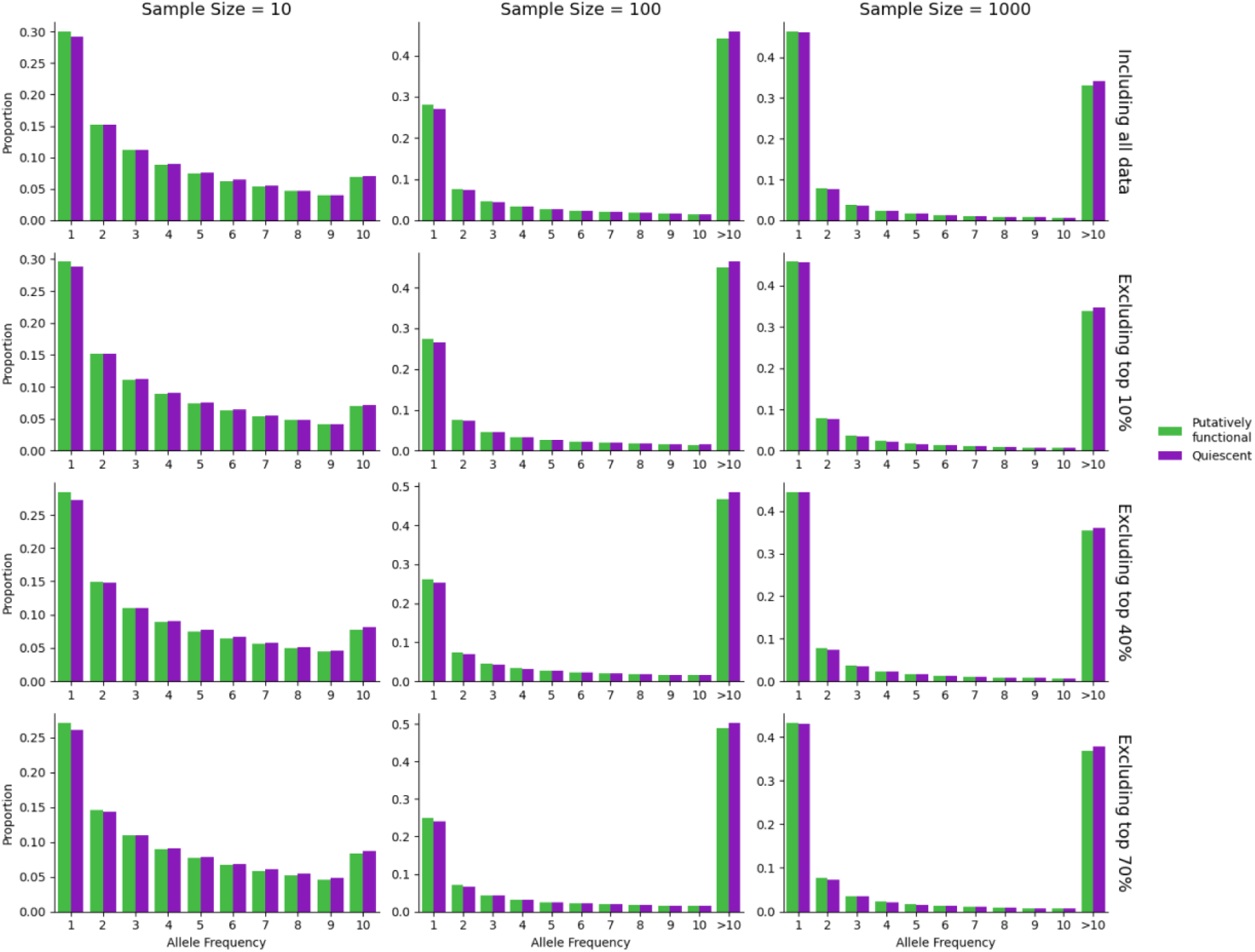
SFSs for putatively functional and quiescent sequence, each subset to 100 Mb total. Each row and column correspond to different sample sizes and PhastCons filtering thresholds, respectively. Sample sizes are in terms of number of haploid genomes. PhastCons filtering excludes sites within the top X% of PhastCons primate-wide or mammalian-wide scores (see Methods). Putatively functional regions overall show a skew towards rare variants and away from common variants.

We used our LRT for homogeneity to formally test whether the SFS for putatively functional sequence differs from that of quiescent sequence. We obtained SFSs for each region type by repeatedly sampling 2 Mb of sequence and then querying SNPs within the sampled sequence. To reduce confounding from LD and background selection, each 2 Mb sample from the putatively functional dataset was matched with a 2 Mb sample from the quiescent dataset such that both samples start within 1000bp of each other. Then, we applied the same bootstrapping procedure we used previously for our simulations. We sampled a subset of SFSs from each of the putatively functional SFSs and the quiescent SFSs, summed each SFS subset, and then applied the LRT for homogeneity. This process was repeated several times, and we used the mean of the resulting LRT statistics to determine statistical significance. We also constructed an empirical null distribution by comparing two sets of quiescent SFSs to each other using the same bootstrap procedure.

LRT results from comparing putatively functional and quiescent variants reveal that the SFSs are remarkably distinct from each other (Figure 6). At the shortest tested sequence length of 100 Mb (first column of Figure 6), all the mean LRT statistics from comparing putatively functional to quiescent exceed their respective significance thresholds across all PhastCons filtering levels. Statistical significance increases with the amount of sequence tested (last three columns of Figure 6). Strikingly, excluding the top 70% of PhastCons sites, the most stringent threshold we tested, still shows a significant difference (last row of Figure 6) in the SFSs between putatively functional regions and quiescent regions. We also observe a stronger signal with decreasing sample size, a trend that echoes the results from our simulations (Supplementary Figure 6). Generally, these results align well with the trends seen in the SFSs (Figure 5).

**Figure 6.**
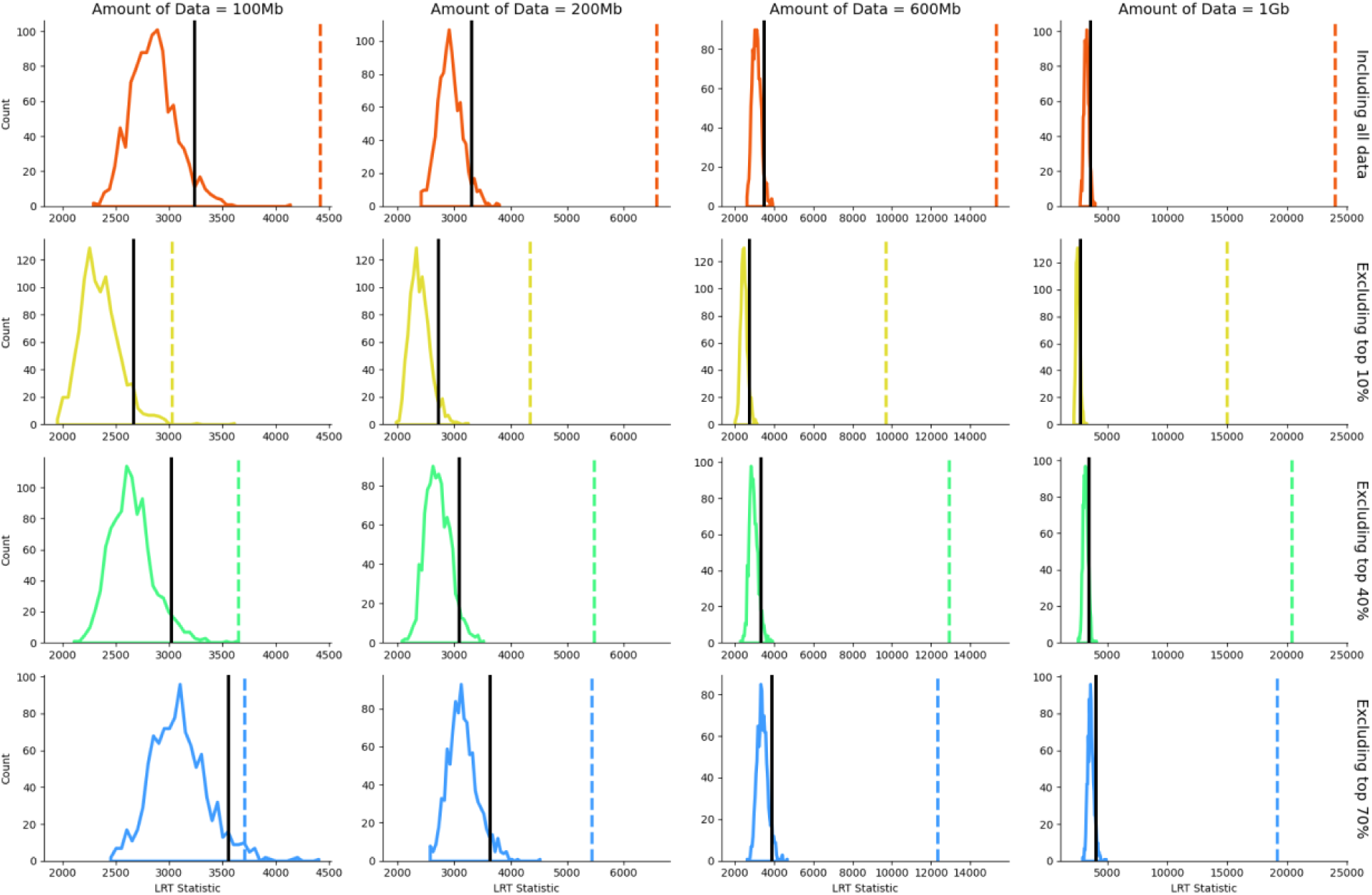
LRT distributions for 1KGP data for sample size of 1000 haploid genomes. Each row and column corresponds to different amounts of tested sequence and PhastCons filtering thresholds, respectively. PhastCons filtering excludes sites within the top X% of PhastCons primate-wide or mammalian-wide scores (see Methods). Empirical null distributions are histograms of LRT statistics generated by comparing quiescent SFSs to each other. Black lines represent the critical values of the respective empirical null distributions. Dashed lines are the mean LRT statistic from comparing putatively functional to quiescent, which are detected as statistically significant across all sample sizes and filtering thresholds shown here.

When just examining the empirical null distributions themselves, we observed a rightward shift as increasing proportions of PhastCons sites are excluded (Supplementary Figure 6). This suggests that SFSs from quiescent regions exhibit greater variance with the progressive removal of conserved sites, leading to larger LRT values in the null distributions. As more PhastCons sites are excluded, the remaining quiescent sequence is, effectively, increasingly enriched for neutrally evolving sites. The Poisson Random Field model predicts that the variance in the number of polymorphisms is greater under neutrality than under negative selection (Sawyer and Hartl 1992), which likely explains this observation. This also implies that even sequences annotated as quiescent may still contain deleterious mutations that are not identified by simple utilization of PhastCons. As truly neutral sites in the LRT for homogeneity would yield a stronger and more accurate signal of negative selection in the putatively functional regions, the detected differences between putatively functional and quiescent SFSs here are likely conservative.

Notably, the magnitude of the differences we observed between quiescent and putatively functional sequences is greater than what we saw in the simulations, even when excluding CNC sites up to the top 70% of PhastCons sites. With this aggressive filtering, we would expect most, if not all, of the variants under negative selection to be removed in both datasets. Yet, our qualitative and quantitative results, i.e. the SFSs and LRT comparisons, respectively, suggest negative selection persists in the putatively functional sequence. Overall, these findings indicate that phylogenetic conservation does not capture all sites where mutations are under negative selection, and the deviations we observe are consistent with cryptic selection driven by functional turnover at putatively functional sites.

### At least 7% of the human genome is under negative selection

We used the 1KGP data in conjunction with simulations to bound the amount of cryptic selection present in the putatively functional regions of the human noncoding genome (Supplementary Figure 7). We processed the 1KGP data as before by separating putatively functional and quiescent variants into two datasets and sampling 2 Mb regions to create SFSs. Unlike in the previous section, we did not downsample the entire putatively functional and quiescent datasets to have equivalent total lengths, as our goal here is not to make rigorous comparisons across PhastCons filtering thresholds. We did exclude varying amounts of conserved sites to assess how conserved sequence affects the detection of cryptic selection. We generated several SFSs using the variants in the two datasets and used our bootstrapping procedure to create empirical null distributions and mean LRT statistics (Supplementary Figure 7B).

We generated new simulations such that downstream analysis would closely match how we processed the 1KGP data. Simulation replicates modeled genomes containing coding, conserved, quiescent, and BGNC regions. We determine the locations for these region types using functional genomic annotations, as we did for our complex, real-world simulations. Each simulation replicate contained at least 2 Mb each of quiescent and BGNC sequence. BGNC regions had either no cryptic selection or 10-80% of the mutations subject to the ncDFE, equivalent to 2.61-20.81% of BGNC mutations under negative selection (Supplementary Table 1). We extracted variants within BGNC and quiescent regions to construct SFSs, focusing only on using a sample size of 1000 haploid genomes (i.e. 500 individuals). Because the number of quiescent and BGNC sites varied across simulation replicates, SFSs were downsampled to represent exactly 2 Mb of sequence for each region type. We applied the same bootstrapping procedure described previously to test the effect of varying sequence lengths (Supplementary Figure 7A). Quiescent SFSs were compared to generate empirical null distributions, and quiescent and BGNC SFSs were compared to generate alternative distributions. Since BGNC regions were simulated under multiple levels of cryptic selection, we performed separate comparisons for each level ranging from 10-80%, and the resulting alternative distributions served as thresholds for estimating the degree of cryptic selection in the 1KGP data. To enable direct comparison between simulations and 1KGP data, all null distributions, alternative distributions, and mean LRT statistics were standardized (see Methods). The standardized 1KGP mean LRT statistics were then evaluated relative to the standardized simulation-based thresholds. The lower and upper thresholds were converted to estimates of genome-wide constraint using Supplementary Table 3.

We find that varying the amounts of PhastCons sites excluded in the putatively functional and quiescent datasets substantially alters our estimates of genome-wide constraint (Figure 7). When both putatively functional and quiescent datasets from the 1KGP undergo the same PhastCons filtering (Figure 7A-B), the mean LRT statistics fall between the 10% and 20% thresholds of the simulation data. This is equivalent to 2.6-5.2% of putatively functional mutations under negative selection (Supplementary Table 1). However, converting the thresholds to amounts of genome-wide negative selection results in different ranges depending on how many PhastCons sites are excluded (Supplementary Table 3). When excluding the top 10% of PhastCons sites in both datasets (Figure 7A), the 1KGP mean LRT statistic, we estimate approximately 7.02-8.39% of the genome under constraint. Excluding more conserved sites, up to the top 20% of PhastCons sites in both datasets (Figure 7B), results in a slightly higher, but fairly similar, estimate of 10.5-11.6% of genome-wide constraint. We also investigated whether excluding PhastCons sites in only the quiescent data creates a greater signal of cryptic selection in the putatively functional data (Figure 7C-D), as expected. When excluding the top 10% of PhastCons sites in just the quiescent data (Figure 7C), we observe that the mean LRT statistic for the 1KGP data falls between the 40% and 60% thresholds, equivalent to 9.31-12.7% of the genome undergoing negative selection. When excluding the top 20% of PhastCons sites in only the quiescent data (Figure 7D), the mean LRT statistic for the 1KGP data falls above the 60% and even exceeds the 80% threshold at a dataset size of 100 Mb, which suggests that more than 17.4% of the genome is under negative selection. Across these different definitions of putatively functional and quiescent, our estimates of genome-wide constraint vary widely, with the lowest estimate around 7% and the highest exceeding 17%.

**Figure 7.**
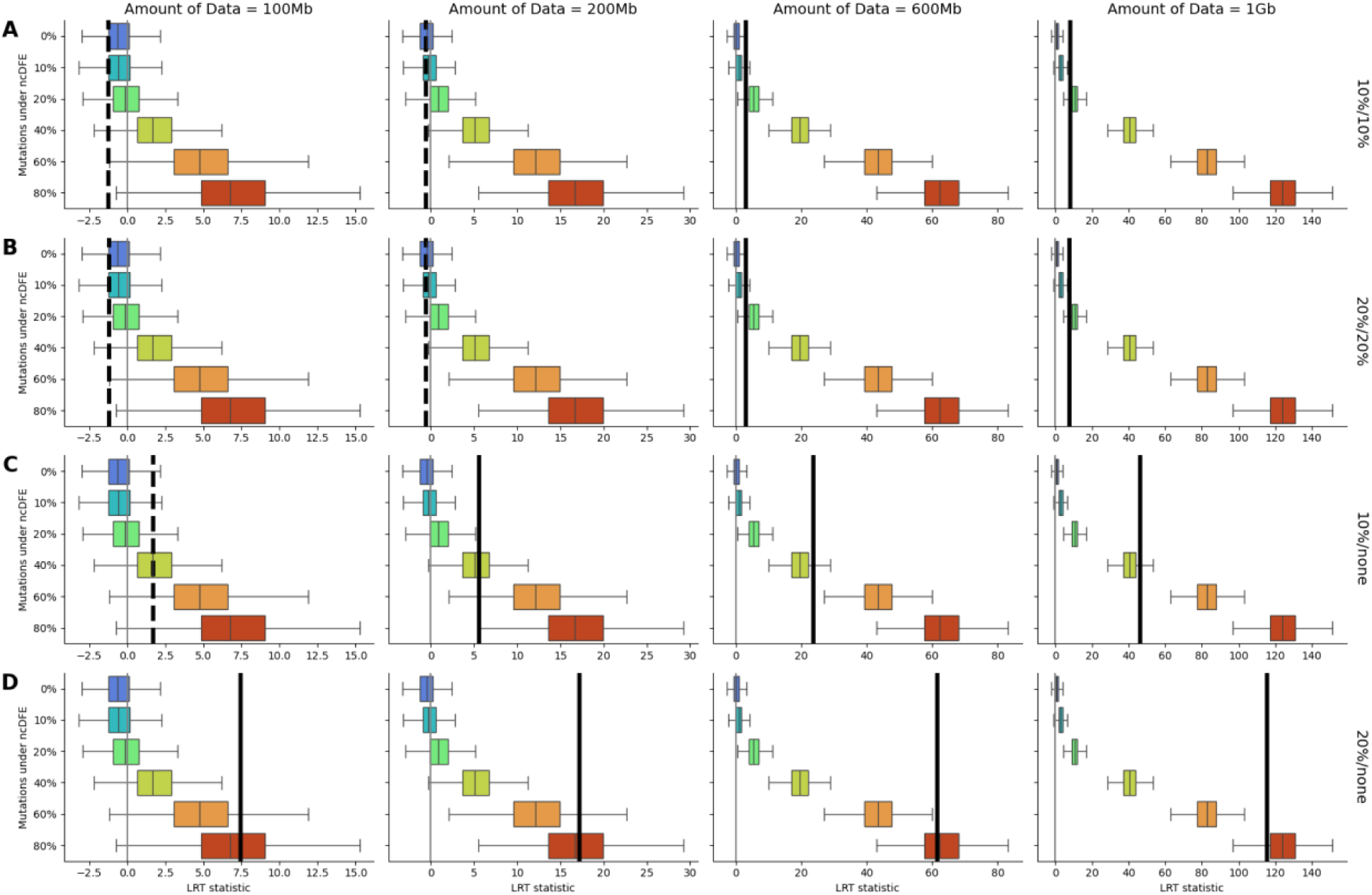
LRT analysis for 1KGP data and matched simulations for a sample size of 1000 haploid genomes. Each column corresponds to a different dataset size, and each row corresponds to a different 1KGP dataset comparison. LRT distributions from BGNC vs. quiescent for simulations (colored boxplots) vary across columns only, while mean LRT statistics from putatively functional vs. quiescent for 1KGP data (black lines) vary across all plots. Solid and dashed lines indicate statistically significant and non-significant outcomes, respectively. All null distributions are standardized to facilitate comparison and are not shown. PhastCons filtering is applied as follows: the top 10% (A) and the top 20% (B) of PhastCons sites are excluded in both the putatively functional and quiescent datasets, while the top 10% (C) and the top 20% (D) of PhastCons sites is excluded only in the quiescent data.

Altogether, we find that method used to define our quiescent data strongly influences our estimate of cryptic selection in the putatively functional data. This sensitivity arises because the quiescent data, which serves as a baseline, is used to measure excess negative selection in the putatively functional data. When we exclude more conserved sites from the quiescent data, the baseline shifts to be more “neutral”-like, thus making the putatively functional sites appear more constrained. When fewer conserved sites are excluded from the quiescent data, these regions retain sites under negative selection, thus comparisons of quiescent to putatively functional results in the latter appearing to be less constrained. Our results corroborate this rationale. When conserved sites are excluded equally from both putatively functional and quiescent sequence, estimates of genome-wide constraint remain similar; however, excluding conserved sites from quiescent sequence alone produces progressively higher constraint estimates as more sites are removed. These patterns emphasize that estimates of constraint are highly dependent on how the baseline, or neutrally evolving, regions and the test, or putatively functional, regions are defined.

Our findings reveal that mutations in at least 7% of sites in the human genome are subjected to negative natural selection. The actual amount is likely much higher, given the large shifts in estimates of constraint based on the filtering of conserved sites in the datasets we compare. Broadly though, what we have observed suggests that negative selection is widespread in non-conserved bases in the human genome, and the true amount of constraint is obscured when the baseline neutral dataset is not truly neutral.

## DISCUSSION

In this work, we investigated the effects of cryptic selection, i.e. the negative selection on mutations in regions of the genome not phylogenetically conserved across species, on patterns of genetic variation. We developed an LRT to compare SFSs with and without cryptic selection, and we used simulations to show that our test can detect as little as 0.781% of mutations affected by cryptic selection (equivalent to 3% of BGNC sites under the ncDFE, see Supplementary Table 1). Successful detection depended on the amount of data and the number of individuals tested. As expected, testing a greater amount of sequence resulted in greater power, but interestingly, we observed the greatest power at an intermediate sample size of 100 individuals. This was likely due to the intrinsic properties of the LRT for homogeneity, as discussed earlier (see Results). We used our LRT for homogeneity to detect cryptic selection in the human noncoding genome by comparing polymorphism data from quiescent and putatively functional sites. We found that signals of cryptic selection are quite pervasive, even when removing the top conserved sites from our analysis. This finding suggests many deleterious mutations are lurking outside of the most phylogenetically conserved regions of the human genome.

Intriguingly, we found that mutations in quiescent regions themselves may also be deleterious, as the null distributions for our LRT built using quiescent SFSs shift to be smaller (i.e. look more “neutral”) as more PhastCons sites are excluded. We also observe this in the joint analysis of simulation and real data. We find that the estimated amount of sites under negative selection in the putatively functional regions varies widely depending on how aggressively we exclude conserved sites from the quiescent regions. These results show that cryptic selection cannot be fully characterized without properly accounting for the negative selection that remains in the “neutral” sequence used as the baseline.

Our observations of pervasive negative selection concur with several studies that find that conservation scores miss many mutations undergoing negative selection. Rands et al. (2014) found an enrichment of ENCODE-identified functional elements within pan-mammalian non-conserved sequences that were identified as constrained by the inference method they developed. Their method also identified constrained sequences that have no functional annotations in ENCODE. Huber et al. (2020) demonstrated that conservation scores can miss both weakly and strongly deleterious mutations. They showed that in simulations, both weakly and strongly deleterious mutations can be assigned the largest constraint score value. Adding functional turnover exacerbates this issue, as they observe that neutrally evolving sites start showing high levels of constraint. Recent work by Di et al. (2025) discovered that noncoding mutations are inferred to be under selection even in some of the least conserved regions of the genome. Specifically, they found that the top 20% of the most conserved sites include only about half of the mutations undergoing negative selection, and the 50% of the genome that is the least constrained has 20% of all deleterious mutations in the genome. These studies all point to functional turnover to explain their results and reinforce the trends we see in our 1KGP data analysis. Overall, our work adds to the increasing bank of evidence that unconstrained sites carry considerable deleterious mutational load.

Why do comparative genomic approaches fail to capture sites under functional turnover? The answer may lie in the evolution of regulatory DNA, which is, in a large part, due to epistasis and genetic drift. Bullaughey (2011) used simulations of enhancer sequences undergoing stabilizing selection over long timescales to investigate how functional turnover may occur. Their results show that mutations that are deleterious in some contexts can be effectively neutral in others, and when in the right context, drift can carry them to fixation. Essentially, the fate of new mutations is context dependent, i.e. epistasis drives changes in the selection landscape of a locus. So, a new mutation may drive functional turnover by itself or may create conditions for other mutations to do so. Because the trajectory of each regulatory sequence is stochastic, different simulation replicates, even under identical selective regimes, diverge in their functional repatterning. Bullaughey argues that each simulation replicate, in a sense, represents an alternative realization of evolutionary history. Thus, effectively neutral sites may appear conserved while functional sites may not. Fixations of effectively neutral mutations may not be directly functional, yet can create opportunities for functional repatterning at other, non-conserved nucleotides. Therefore, conservation does not necessarily imply functionality, and lack of conservation does not preclude it. These functional sites experience negative selection and population genetic summary statistics such as the SFS can capture these perturbations while conservation-based metrics cannot. Thus, focusing on patterns of genetic variation may better illuminate functionality in the human genome.

Using our SFS-based comparisons, we estimate that mutations in at least 7% of the genome are under negative selection. Several prior studies have estimated that when including functional turnover, 8-15% of the human genome is under evolutionary constraint (Asthana, Roytberg, et al. 2007; Di et al. 2025; Meader et al. 2010; Ponting and Hardison 2011; The ENCODE Project Consortium et al. 2020; Ward and Kellis 2012). Unlike these works, our approach is not designed to produce a precise genome-wide estimate of constraint. Instead, our goal is to detect deviations in the SFS attributable specifically to cryptic selection. Yet, it is still reassuring that the amount of selection required to generate the difference between the quiescent and putatively functional regions that we detected is in line with previous estimates. A strength of our back-of-the-envelope estimate is that our approach makes very few underlying assumptions. Rather than modeling variation over time or using other inference frameworks to estimate selection, we simply rely on direct comparisons of SFSs using our LRT for homogeneity.

Our analyses suggest that the proportion of sites under selective constraint could exceed 17% depending on how our control, i.e. neutral sequence, is defined (Figure 7). This appears to conflict with the theoretical maximum of 15% functional sites proposed by Graur (2017). By using the Crow definition of mutational load, in which selection acts on absolute fitness relative to a deleterious mutation-free individual, a functional fraction exceeding 15% of the genome would require an unrealistically high number of offspring per individual to maintain a constant population size. However, Lesecque et al. (2012) argued that the more biologically appropriate Wallace definition of mutational load should be used instead, in which selection acts on the variation in fitness in a population due to competition between individuals. Under this relative fitness framework, high levels of deleterious mutations are compatible with realistic reproductive rates, even across a range of selection strengths and reproductive strategies. Our estimates of constraint are therefore not biologically implausible.

Our work emphasizes that conservation scores are an imperfect metric for identifying putatively selected sites, as we find that cryptic selection is pervasive in the noncoding genome. This has implications for studies conducting inference using patterns of genetic variation. Inference of human demographic history requires putatively neutral sites, yet some studies use intergenic SNPs with little to no further filtering (Gravel et al. 2011; Gutenkunst et al. 2009). Our findings that in these regions, more mutations than previously appreciated may be deleterious could indicate that current estimates of demographic history are erroneously inferring stronger periods of recent population growth. Estimating the strength of background selection also requires identifying neutral sites on which background selection acts on. Some of these studies use conservation scores to aid in identifying putatively selected and putatively neutral sites (Buffalo and Kern 2024; McVicker et al. 2009; Phung et al. 2016). However, these putatively neutral sites may exhibit signals of reduced diversity or divergence simply due to the direct effects, rather than the indirect effects, of selection. Likewise, putatively selected sites identified by conservation scores may include neutrally evolving sites. Thus, the strength of background selection may be overestimated in these studies.

Our conclusions come with some important caveats. First, we rely on functional annotations to define quiescent regions. However, these annotations are imperfect, as our analyses make clear. Regions with very little transcriptional activity are inferred to be quiescent by ChromHMM (Hoffman et al. 2013). We use these annotations and further filter out ENCODE-identified enhancers and promoters to enrich for neutrally evolving sites. Nonetheless, we find that negative selection persists in these quiescent regions, as excluding conserved sites makes quiescent SFSs more neutral-like and alters the signal of cryptic selection in putatively functional regions. These findings are perhaps unsurprising, as the regions annotated as quiescent by ChromHMM may in fact be functional in other cell types or under other biological conditions, and still subjected to negative selection (Grujic et al. 2020; Hoffman et al. 2013). Nevertheless, this limitation is intrinsic to the functional genomic data rather than our methodology and is part of the phenomenon that we highlight in this work, i.e. negative selection and neutrality are not cleanly separable in the human genome. As a result, our estimates of cryptic selection are likely conservative due to this caveat. Second, we attribute differences between putatively functional and quiescent regions to the effects of negative selection, as the SFSs exhibit the characteristic pattern of increased rare variants and decreased common variants (Figure 5). However, this skew may be caused by relatively lower mutation rates within putatively functional regions, which may reduce genetic diversity and inflate differences detected in the SFS.

Altogether, our findings support the view that functional turnover plays an important role in shaping patterns of genetic variation in the human genome. Importantly, our results demonstrate that population genetic approaches can illuminate aspects of genomic constraint that comparative genomic methods cannot. The SFS is a simple yet powerful statistic that is sensitive to the effects of selection in genetic variation data, and our work offers a complementary approach to determine the extent of functionality in the human genome. More broadly, we argue that future work in population genetics and molecular evolution should account for functional turnover when defining neutral baselines, interpreting patterns of diversity, and estimating the strength of natural selection. Failing to do so risks systematically missing key regions under selective constraint in the human genome, and in turn, obscuring our understanding of how natural selection has shaped human evolution.

## METHODS

### Human genome annotations

We obtained two sets of PhastCons scores for each site in the human genome from the UCSC Genome Browser: one calculated from alignments across 17 primate species and one from alignments across 470 mammalian species (Casper et al. 2025; Siepel et al. 2005). Sites were binned by PhastCons score percentile as per Di et al. (2025). We combined primate-wide and mammalian-wide score bins such that any estimates from our analyses were as conservative as possible. Quiescent and heterochromatin annotations were acquired from ChromHMM (Ernst and Kellis 2012; Vu and Ernst 2022). Heterochromatin and all ENCODE-annotated human enhancer and promoter sites (The ENCODE Project Consortium et al. 2020) were excluded from the quiescent annotations. Gap and centromere coordinates were retrieved from the UCSC Genome Browser (Casper et al. 2025).

We used various functional genomic annotations to build the genome structures for our complex and 1KGP-matched simulations. For coding elements, we used CDS annotations obtained from Ensembl/Havana Release 104 via the stdpopsim Python library (Adrion et al. 2020). We used the sites within the top 5% of PhastCons scores for CNC elements in the complex, real-world simulations and the top 10% for the 1KGP-matched simulations. We used quiescent annotations in only the 1KGP-matched simulations. Per-bp recombination and mutation rate maps were obtained from HapMap Phase II via stdpopsim (Adrion et al. 2020). Any sequence not labeled as coding, CNC, or quiescent was labeled as BGNC. Any genomic regions that spanned gaps or centromeres were not used to create genome structures for our simulations.

We used functional genomic annotations to also filter for SNPs within putatively functional and quiescent regions from the 1KGP data. CDS annotations used for filtering SNPs were obtained from Gencode Release 45 Version GRCh38 (Frankish et al. 2023). We used varying PhastCons thresholds to identify and exclude CNC SNPs. Any SNPs not categorized as quiescent, heterochromatin, CDS, and CNC were categorized as putatively functional.

### Simulations

We performed forward-in-time Wright-Fisher simulations using SLiM version 4.0.1 (Haller and Messer 2023) and modeled European demographic history estimated by Kim et al. (2017). First, we performed simple simulations with synthetic genome structures. Each simulation replicate modeled an approximately 20 Mb chromosome with mutations occurring at a rate of 2e-8 per bp per generation (Rodrigues et al. 2024). We considered two different chromosomal structures. First, chromosome structures were randomly generated with a fixed number of equally sized coding and noncoding regions. Here, the chromosome consisted of 1000 coding segments (each 300bp), 2000 CDS-flanking CNC segments (each 500bp), 1002 BGNC segments (each 18500bp), and one 1 Mb large neutrally evolving segment. We set a constant recombination rate of 1e-8 per bp per generation for the entire chromosome. Then, we created another dataset of simulations with realistic genome structures by integrating publicly available genomic annotations. Here, each chromosome was created by sampling a 20 Mb segment without unmappable regions (i.e. gaps and centromeres) from the human genome. We labeled sites as CDS, CNC, or BGNC based on human functional genomic annotations (see above section).

Coding regions accumulated deleterious and neutral mutations at a ratio of 2.31:1, and selection coefficients for deleterious mutations were drawn from a gamma DFE with parameters estimated by Kim et al. (2017). Selection coefficients for mutations in CNC and BGNC regions were drawn from a DFE inferred from ENCODE candidate cis-regulatory regions (ncDFE) (Torgerson et al. 2009). BGNC regions were simulated to have 0%, 1%, 3%, or 5% of mutations with selection coefficients drawn from the ncDFE to emulate the effects of cryptic selection. See Supplementary Table 1 for the actual proportion of deleterious mutations in each of these scenarios. We ran 100 simulation replicates for each cryptic selection condition, and each set of replicates used the same set of 100 randomly generated chromosome structures. We ran 100 extra simulation replicates with different chromosome structures for 0% to facilitate construction of an empirical null distribution. Each simulation replicate output SFSs for sample sizes of 10, 100, and 1000 haploid genomes for every type of region simulated, i.e. CDS, CNC, BGNC, and neutral (if applicable).

To quantify the effects of cryptic selection on the SFS, we applied the LRT for homogeneity we developed to SFSs from simulations without cryptic selection and the SFSs from simulations with cryptic selection (Figure 2). Here, we employed a bootstrap procedure. We randomly sampled a subset of SFS from the set of SFSs without cryptic selection and from the set of SFSs with cryptic selection. We summed the SFSs in each subset across SFS bins and then calculated the LRT statistic using the two aggregate SFSs. This process was iterated 1000 times to produce 1000 LRT statistics, and these were averaged to produce one mean LRT statistic. Since the total amount of sequence considered may have an effect on power to detect cryptic selection, we varied the size of the subsets from each of the two simulation sets compared. Subset sizes ranged from 5-50 simulations, equivalent to 100 Mb - 1 Gb of simulated sequence. We obtained mean LRT statistics for each subset size, sample size (10, 100, and 1000 haploid genomes), and strength of cryptic selection (1%, 3%, and 5%).

To create an empirical null distribution, we split 200 simulation replicates with 0% under the ncDFE into two sets of 100 simulations each. We performed the same bootstrap procedure as described above and the resulting 1000 LRT statistics comprised the null distribution. The set of mean LRT statistics derived from comparing simulations with and without cryptic selection were evaluated along with the null distribution with a significance threshold of 0.05 to determine whether the LRT means were significantly different from the empirical null distribution.

### 1KGP Data

We obtained biallelic SNPs from 525 unrelated European individuals from the 1KGP Phase III dataset (Byrska-Bishop et al. 2022) that were within the 1KGP Strict Mask (Auton et al. 2015). SNPs that did not contain “PASS” in the “FILTER” column or that had any missing genotypes were excluded.

We compared SFSs constructed from SNPs in quiescent regions to those from putatively functional regions. For both categories, we removed sites with PhastCons scores above a specified percentile threshold, which we varied from the top 10% to the top 70%, to test the effect of excluding conserved sites and to reduce any residual negative selection from mutations at constrained sites that could obscure differences in the SFS attributable to cryptic selection alone. PhastCons filtering reduces the amount of testable sequence–higher percentile thresholds mean more sites are excluded.

There are fewer quiescent sites than there are putatively functional sites in the genome, so we controlled for this imbalance by downsampling quiescent and putatively functional annotations to 100 Mb across all PhastCons thresholds. To assess whether downsampling introduces any differential LD, we calculated pairwise LD between SNPs in both 100 Mb subsets using PLINK v1.07 (Chang et al. 2015). We limited SNP pairs to only pairs where both SNPs resided in quiescent or both resided in putatively functional regions. We retained SNPs with a minor allele frequency greater than 1% and SNP pairs within 80 Mb of each other. We then computed the number of SNP pairs and mean r^2^ for each of 50 log-spaced pairwise-distance bins for each annotation type. We also computed cumulative distributions of r^2^ values, restricted to SNP pairs with pairwise distance within 1 kb - 1 Mb.

We constructed SFSs from quiescent and putatively functional SNPs for sample sizes of 10, 100, and 1000 haploid genomes (i.e. 5, 50, and 500 individuals). To control for local linkage effects, we randomly sampled 2 Mb of quiescent and putatively functional sequence from matched genomic locations, with starting positions within 1000 bp of each other. We repeated this sampling 1000 times to yield 1000 paired SFSs for each region type. We created an additional set of 1000 quiescent SFSs for the empirical null distribution.

We compared quiescent and putatively functional SFSs using the LRT for homogeneity and used bootstrapping to test the effects of varying amounts of sequence on the detection of cryptic selection. The entire process closely mirrors our treatment of the simulations. Variable-sized subsets of SFSs were randomly sampled from each of our quiescent and putatively functional SFS sets. Subset sizes ranged from 50 - 500 SFSs, equivalent to 100 Mb - 1 Gb of sequence. The SFSs in each subset were summed and compared using the LRT for homogeneity. We iterated this bootstrapping 1000 times and averaged the resulting 1000 LRT statistics to produce one mean LRT statistic. We repeated this process for all three sample sizes. We also applied this same procedure to two sets of quiescent SFSs to create an empirical null distribution for all SFS subset sizes and sample sizes. Using a significance threshold of 0.05, we evaluated whether the mean LRT statistics from comparing quiescent and putatively functional SFSs were significantly deviated from their respective empirical null distributions.

### Estimating constraint in the human genome

We designed simulations to match our comparison of putatively functional and quiescent SNPs from the 1KGP (Supplementary Figure 7). We sampled 1000 genomic regions with integrated CDS, CNC, and quiescent annotations. Sites in the top 10% of PhastCons scores were treated as CNC, and those outside of CDS, CNC, and quiescent were treated as BGNC. We ensured that each sample contained at least 2 Mb of quiescent and BGNC sequence each to reflect the sampling scheme used for the 1KGP data (see below) but also capped sampled region lengths at 8 Mb for computational tractability and efficiency. Selection coefficients for mutations introduced in CDS, CNC, and BGNC regions used the same DFE parameters as in previous simulations. Quiescent regions accumulated only neutral mutations. BGNC regions were simulated under several conditions: 0%, 10%, 20%, 40%, 60%, or 80% of mutations with selection coefficients drawn from the ncDFE (see Supplementary Table 1 for the corresponding proportions of BGNC mutations under negative selection).

As in previous simulations, we generated SFSs for sample sizes of 10, 100, and 1000 haploid genomes for all genomic region types for all simulation replicates. Although simulated genomic regions were sampled to have at least 2 Mb of both quiescent and BGNC sequence, the actual lengths of each type varied widely among the simulation replicates, ranging from 2 Mb to roughly 5 Mb. To control for this variability, we downsampled the SFSs to represent exactly 2 Mb of each type of sequence. The simulations were randomly split into two sets of 500 each (Supplementary Figure 7A). Quiescent SFSs from each set of 500 replicates were compared to each other using the LRT for homogeneity to create an empirical null distribution. For the alternative distribution, we randomly split the simulations again into two sets of 500 each and compared quiescent SFSs and putatively functional SFSs using the LRT for homogeneity. We treated our 1KGP data same as before, with some modifications to better reflect the simulation scheme we used. We excluded either the top 10% or top 20% of PhastCons sites from both putatively functional and quiescent datasets, or from the quiescent dataset alone. Here, we sampled 2 Mb each of quiescent and putatively functional sequence without restricting the two samples to start within 1000bp of each other, as was done in the previous analysis. This is because the previous analysis compared quiescent and putatively functional sequences directly against each other across PhastCons thresholds, whereas here we compare 1KGP data against simulations, which are divided into independent, equally sized sets. Accordingly, the two 2 Mb samples are treated as independent. We restricted the total genomic window encompassing both samples to at most 8 Mb, consistent with the genome sizes used in the simulations. SNPs within each 2 Mb sample were used to construct SFSs, and this sampling was repeated to generate two sets of 500 quiescent SFSs and one set of 500 putatively functional SFSs. The two quiescent SFS sets were used to construct empirical null distributions, while one quiescent and one putatively functional SFS set were compared using the LRT for homogeneity to construct alternative distributions and obtain the mean LRT statistics (Supplementary Figure 7B).

Although our simulations were modeled to closely fit our 1KGP dataset sampling scheme, simulation empirical null distributions did not match 1KGP empirical null distributions. To adjust for this difference, we standardized all empirical null distributions to become Z-scores for ease of comparison (Supplementary Figure 7). For both the simulation and 1KGP data, we used the means and standard deviations of the null distributions to standardize the alternative distributions. Alternative distributions from the simulation data served as thresholds to determine the level of constraint in the 1KGP data. We determined the lower and upper bounds of genome-wide constraint by converting the lower and upper simulation data thresholds using Supplementary Table 3, which was computed by multiplying the proportion of sites under selection stipulated by the DFEs for each type of sequence, i.e. coding, putatively functional, conserved, and quiescent, with their total length in the genome.

## ACKNOWLEDGEMENTS

This work was supported by the National Institutes of Health grant R35GM119856 to KEL.

## REFERENCE

Adrion, Jeffrey R, Christopher B Cole, Noah Dukler, et al. 2020. “A Community-Maintained Standard Library of Population Genetic Models.” eLife 9 (June): e54967. 10.7554/eLife.54967.

Arneson, Adriana, and Jason Ernst. 2019. “Systematic Discovery of Conservation States for Single-Nucleotide Annotation of the Human Genome.” Communications Biology 2 (1): 248. 10.1038/s42003-019-0488-1.

Asthana, Saurabh, William S. Noble, Gregory Kryukov, Charles E. Grant, Shamil Sunyaev, and John A. Stamatoyannopoulos. 2007. “Widely Distributed Noncoding Purifying Selection in the Human Genome.” Proceedings of the National Academy of Sciences 104 (30): 12410–15. 10.1073/pnas.0705140104.

Asthana, Saurabh, Mikhail Roytberg, John Stamatoyannopoulos, and Shamil Sunyaev. 2007. “Analysis of Sequence Conservation at Nucleotide Resolution.” PLoS Computational Biology 3 (12): e254. 10.1371/journal.pcbi.0030254.

Auton, Adam, Gonçalo R. Abecasis, David M. Altshuler, et al. 2015. “A Global Reference for Human Genetic Variation.” Nature 526 (7571): 68–74. 10.1038/nature15393.

Bejerano, Gill, Michael Pheasant, Igor Makunin, et al. 2004. “Ultraconserved Elements in the Human Genome.” Science 304 (5675): 1321–25. 10.1126/science.1098119.

Birney, Ewan, John A. Stamatoyannopoulos, Anindya Dutta, et al. 2007. “Identification and Analysis of Functional Elements in 1% of the Human Genome by the ENCODE Pilot Project.” Nature 447 (7146): 799–816. 10.1038/nature05874.

Boffelli, Dario, Jon McAuliffe, Dmitriy Ovcharenko, et al. 2003. “Phylogenetic Shadowing of Primate Sequences to Find Functional Regions of the Human Genome.” Science 299 (5611): 1391–94. 10.1126/science.1081331.

Buffalo, Vince, and Andrew D. Kern. 2024. “A Quantitative Genetic Model of Background Selection in Humans.” PLOS Genetics 20 (3): e1011144. 10.1371/journal.pgen.1011144.

Bullaughey, Kevin. 2011. “Changes in Selective Effects Over Time Facilitate Turnover of Enhancer Sequences.” Genetics 187 (2): 567–82. 10.1534/genetics.110.121590.

Bustamante, Carlos D, John Wakeley, Stanley Sawyer, and Daniel L Hartl. 2001. “Directional Selection and the Site-Frequency Spectrum.” Genetics 159 (4): 1779–88. 10.1093/genetics/159.4.1779.

Byrska-Bishop, Marta, Uday S. Evani, Xuefang Zhao, et al. 2022. “High-Coverage Whole-Genome Sequencing of the Expanded 1000 Genomes Project Cohort Including 602 Trios.” Cell 185 (18): 3426–3440.e19. 10.1016/j.cell.2022.08.004.

Casper, Jonathan, Matthew L Speir, Brian J Raney, et al. 2025. “The UCSC Genome Browser Database: 2026 Update.” Nucleic Acids Research 54 (D1): D1331–35. 10.1093/nar/gkaf1250.

Chang, Christopher C, Carson C Chow, Laurent CAM Tellier, Shashaank Vattikuti, Shaun M Purcell, and James J Lee. 2015. “Second-Generation PLINK: Rising to the Challenge of Larger and Richer Datasets.” GigaScience 4 (1): s13742-015-0047–0048. 10.1186/s13742-015-0047-8.

Charlesworth, Brian, and Jeffrey D. Jensen. 2021. “Effects of Selection at Linked Sites on Patterns of Genetic Variability.” Annual Review of Ecology, Evolution, and Systematics 52 (Volume 52, 2021): 177–97. 10.1146/annurev-ecolsys-010621-044528.

Chiaromonte, F., R.J. Weber, K.M. Roskin, M. Diekhans, W.J. Kent, and D. Haussler. 2003. “The Share of Human Genomic DNA under Selection Estimated from Human-Mouse Genomic Alignments.” Cold Spring Harbor Symposia on Quantitative Biology 68 (0): 245–54. 10.1101/sqb.2003.68.245.

Christmas, Matthew J., Irene M. Kaplow, Diane P. Genereux, et al. 2023. “Evolutionary Constraint and Innovation across Hundreds of Placental Mammals.” Science (New York, N.Y.) 380 (6643): eabn3943. 10.1126/science.abn3943.

Cooper, Gregory M., Michael Brudno, Eric A. Stone, Inna Dubchak, Serafim Batzoglou, and Arend Sidow. 2004. “Characterization of Evolutionary Rates and Constraints in Three Mammalian Genomes.” Genome Research 14 (4): 539–48. 10.1101/gr.2034704.

Cooper, Gregory M., Eric A. Stone, George Asimenos, Eric D. Green, Serafim Batzoglou, and Arend Sidow. 2005. “Distribution and Intensity of Constraint in Mammalian Genomic Sequence.” Genome Research 15 (7): 901–13. 10.1101/gr.3577405.

Dermitzakis, Emmanouil T., and Andrew G. Clark. 2002. “Evolution of Transcription Factor Binding Sites in Mammalian Gene Regulatory Regions: Conservation and Turnover.” Molecular Biology and Evolution 19 (7): 1114–21. 10.1093/oxfordjournals.molbev.a004169.

Dermitzakis, Emmanouil T., Alexandre Reymond, Nathalie Scamuffa, et al. 2003. “Evolutionary Discrimination of Mammalian Conserved Non-Genic Sequences (CNGs).” Science 302 (5647): 1033–35. 10.1126/science.1087047.

Di, Chenlu, Swetha Ramesh, Jason Ernst, and Kirk E. Lohmueller. 2025. “The Landscape of Fitness Effects of Putatively Functional Noncoding Mutations in Humans.” Preprint, bioRxiv, May 14. 10.1101/2025.05.14.654124.

Doolittle, W. Ford. 2013. “Is Junk DNA Bunk? A Critique of ENCODE.” Proceedings of the National Academy of Sciences 110 (14): 5294–300. 10.1073/pnas.1221376110.

Ernst, Jason, and Manolis Kellis. 2012. “ChromHMM: Automating Chromatin-State Discovery and Characterization.” Nature Methods 9 (3): 215–16. 10.1038/nmeth.1906.

Excoffier, Laurent, Isabelle Dupanloup, Emilia Huerta-Sánchez, Vitor C. Sousa, and Matthieu Foll. 2013. “Robust Demographic Inference from Genomic and SNP Data.” PLoS Genetics 9 (10): e1003905. 10.1371/journal.pgen.1003905.

Frankish, Adam, Sílvia Carbonell-Sala, Mark Diekhans, et al. 2023. “GENCODE: Reference Annotation for the Human and Mouse Genomes in 2023.” Nucleic Acids Research 51 (D1): D942–49. 10.1093/nar/gkac1071.

Garber, Manuel, Mitchell Guttman, Michele Clamp, Michael C. Zody, Nir Friedman, and Xiaohui Xie. 2009. “Identifying Novel Constrained Elements by Exploiting Biased Substitution Patterns.” Bioinformatics 25 (12): i54–62. 10.1093/bioinformatics/btp190.

Gazave, Elodie, Diana Chang, Andrew G. Clark, and Alon Keinan. 2013. “Population Growth Inflates the Per-Individual Number of Deleterious Mutations and Reduces Their Mean Effect.” Genetics 195 (3): 969–78. 10.1534/genetics.113.153973.

Graur, D., Y. Zheng, N. Price, R. B. R. Azevedo, R. A. Zufall, and E. Elhaik. 2013. “On the Immortality of Television Sets: ‘Function’ in the Human Genome According to the Evolution-Free Gospel of ENCODE.” Genome Biology and Evolution 5 (3): 578–90. 10.1093/gbe/evt028.

Graur, Dan. 2017. “An Upper Limit on the Functional Fraction of the Human Genome.” Genome Biology and Evolution 9 (7): 1880–85. 10.1093/gbe/evx121.

Gravel, Simon, Brenna M. Henn, Ryan N. Gutenkunst, et al. 2011. “Demographic History and Rare Allele Sharing among Human Populations.” Proceedings of the National Academy of Sciences 108 (29): 11983–88. 10.1073/pnas.1019276108.

Grujic, Olivera, Tanya N. Phung, Soo Bin Kwon, et al. 2020. “Identification and Characterization of Constrained Non-Exonic Bases Lacking Predictive Epigenomic and Transcription Factor Binding Annotations.” Nature Communications 11 (1): 6168. 10.1038/s41467-020-19962-9.

Gutenkunst, Ryan N., Ryan D. Hernandez, Scott H. Williamson, and Carlos D. Bustamante. 2009. “Inferring the Joint Demographic History of Multiple Populations from Multidimensional SNP Frequency Data.” PLoS Genetics 5 (10): e1000695. 10.1371/journal.pgen.1000695.

Haller, Benjamin C., and Philipp W. Messer. 2023. “SLiM 4: Multispecies Eco-Evolutionary Modeling.” The American Naturalist 201 (5): E127–39. 10.1086/723601.

Hoffman, Michael M., Jason Ernst, Steven P. Wilder, et al. 2013. “Integrative Annotation of Chromatin Elements from ENCODE Data.” Nucleic Acids Research 41 (2): 827–41. 10.1093/nar/gks1284.

Huber, Christian D., Bernard Y. Kim, and Kirk E. Lohmueller. 2020. “Population Genetic Models of GERP Scores Suggest Pervasive Turnover of Constrained Sites across Mammalian Evolution.” PLOS Genetics 16 (5): e1008827. 10.1371/journal.pgen.1008827.

Kallenberg, W. C. M., J. Oosterhoff, and B. F. Schriever. 1985. “The Number of Classes in Chi-Squared Goodness-of-Fit Tests.” Journal of the American Statistical Association 80 (392): 959–68. 10.1080/01621459.1985.10478211.

Kellis, Manolis, Barbara Wold, Michael P. Snyder, et al. 2014. “Defining Functional DNA Elements in the Human Genome.” Proceedings of the National Academy of Sciences 111 (17): 6131–38. 10.1073/pnas.1318948111.

Kim, Bernard Y, Christian D Huber, and Kirk E Lohmueller. 2017. “Inference of the Distribution of Selection Coefficients for New Nonsynonymous Mutations Using Large Samples.” Genetics 206 (1): 345–61. 10.1534/genetics.116.197145.

King, David C., James Taylor, Laura Elnitski, Francesca Chiaromonte, Webb Miller, and Ross C. Hardison. 2005. “Evaluation of Regulatory Potential and Conservation Scores for Detecting Cis-Regulatory Modules in Aligned Mammalian Genome Sequences.” Genome Research 15 (8): 1051–60. 10.1101/gr.3642605.

Kirkness, Ewen F., Vineet Bafna, Aaron L. Halpern, et al. 2003. “The Dog Genome: Survey Sequencing and Comparative Analysis.” Science 301 (5641): 1898–903. 10.1126/science.1086432.

Lesecque, Yann, Peter D. Keightley, and Adam Eyre-Walker. 2012. “A Resolution of the Mutation Load Paradox in Humans.” Genetics 191 (4): 1321–30. 10.1534/genetics.112.140343.

Lindblad-Toh, Kerstin, Manuel Garber, Or Zuk, et al. 2011. “A High-Resolution Map of Human Evolutionary Constraint Using 29 Mammals.” Nature 478 (7370): 476–82. 10.1038/nature10530.

Margulies, Elliott H., Mathieu Blanchette, NISC Comparative Sequencing Program, David Haussler, and Eric D. Green. 2003. “Identification and Characterization of Multi-Species Conserved Sequences.” Genome Research 13 (12): 2507–18. 10.1101/gr.1602203.

McVicker, Graham, David Gordon, Colleen Davis, and Phil Green. 2009. “Widespread Genomic Signatures of Natural Selection in Hominid Evolution.” PLOS Genetics 5 (5): e1000471. 10.1371/journal.pgen.1000471.

Meader, Stephen, Chris P. Ponting, and Gerton Lunter. 2010. “Massive Turnover of Functional Sequence in Human and Other Mammalian Genomes.” Genome Research 20 (10): 1335–43. 10.1101/gr.108795.110.

Miller, Webb, Kateryna D. Makova, Anton Nekrutenko, and Ross C. Hardison. 2004. “COMPARATIVE GENOMICS.” Annual Review of Genomics and Human Genetics 5 (1): 15–56. 10.1146/annurev.genom.5.061903.180057.

Mouse Genome Sequencing Consortium. 2002. “Initial Sequencing and Comparative Analysis of the Mouse Genome.” Nature 420 (6915): 520–62. 10.1038/nature01262.

Parker, Stephen C. J., Loren Hansen, Hatice Ozel Abaan, Thomas D. Tullius, and Elliott H. Margulies. 2009. “Local DNA Topography Correlates with Functional Noncoding Regions of the Human Genome.” Science (New York, N.Y.) 324 (5925): 389–92. 10.1126/science.1169050.

Phung, Tanya N., Christian D. Huber, and Kirk E. Lohmueller. 2016. “Determining the Effect of Natural Selection on Linked Neutral Divergence across Species.” PLOS Genetics 12 (8): e1006199. 10.1371/journal.pgen.1006199.

Pollard, Katherine S., Melissa J. Hubisz, Kate R. Rosenbloom, and Adam Siepel. 2010. “Detection of Nonneutral Substitution Rates on Mammalian Phylogenies.” Genome Research 20 (1): 110–21. 10.1101/gr.097857.109.

Ponting, Chris P., and Ross C. Hardison. 2011. “What Fraction of the Human Genome Is Functional?” Genome Research 21 (11): 1769–76. 10.1101/gr.116814.110.

Rands, Chris M., Stephen Meader, Chris P. Ponting, and Gerton Lunter. 2014. “8.2% of the Human Genome Is Constrained: Variation in Rates of Turnover across Functional Element Classes in the Human Lineage.” PLoS Genetics 10 (7): e1004525. 10.1371/journal.pgen.1004525.

Reanney, D. 1976. “Extrachromosomal Elements as Possible Agents of Adaptation and Development.” Bacteriological Reviews 40 (3): 552–90. 10.1128/br.40.3.552-590.1976.

Rodrigues, Murillo F, Andrew D Kern, and Peter L Ralph. 2024. “Shared Evolutionary Processes Shape Landscapes of Genomic Variation in the Great Apes.” Genetics 226 (4): iyae006. 10.1093/genetics/iyae006.

Sawyer, S A, and D L Hartl. 1992. “Population Genetics of Polymorphism and Divergence.” Genetics 132 (4): 1161–76. 10.1093/genetics/132.4.1161.

Shabalina, S. 2001. “Selective Constraint in Intergenic Regions of Human and Mouse Genomes.” Trends in Genetics 17 (7): 373–76. 10.1016/S0168-9525(01)02344-7.

Siepel, Adam, Gill Bejerano, Jakob S. Pedersen, et al. 2005. “Evolutionarily Conserved Elements in Vertebrate, Insect, Worm, and Yeast Genomes.” Genome Research 15 (8): 1034–50. 10.1101/gr.3715005.

Smith, Nick G.C., Mikael Brandström, and Hans Ellegren. 2004. “Evidence for Turnover of Functional Noncoding DNA in Mammalian Genome Evolution.” Genomics 84 (5): 806–13. 10.1016/j.ygeno.2004.07.012.

The ENCODE Project Consortium. 2012. “An Integrated Encyclopedia of DNA Elements in the Human Genome.” Nature 489 (7414): 57–74. 10.1038/nature11247.

The ENCODE Project Consortium, Federico Abascal, Reyes Acosta, et al. 2020. “Expanded Encyclopaedias of DNA Elements in the Human and Mouse Genomes.” Nature 583 (7818): 699–710. 10.1038/s41586-020-2493-4.

Torgerson, Dara G., Adam R. Boyko, Ryan D. Hernandez, et al. 2009. “Evolutionary Processes Acting on Candidate Cis-Regulatory Regions in Humans Inferred from Patterns of Polymorphism and Divergence.” PLOS Genetics 5 (8): e1000592. 10.1371/journal.pgen.1000592.

Vu, Ha, and Jason Ernst. 2022. “Universal Annotation of the Human Genome through Integration of over a Thousand Epigenomic Datasets.” Genome Biology 23 (January): 9. 10.1186/s13059-021-02572-z.

Ward, Lucas D., and Manolis Kellis. 2012. “Evidence of Abundant Purifying Selection in Humans for Recently Acquired Regulatory Functions.” Science 337 (6102): 1675–78. 10.1126/science.1225057.

Watterson, G. A. 1975. “On the Number of Segregating Sites in Genetical Models without Recombination.” Theoretical Population Biology 7 (2): 256–76. 10.1016/0040-5809(75)90020-9.

Young, Robert S. 2016. “Lineage-Specific Genomics: Frequent Birth and Death in the Human Genome.” BioEssays 38 (7): 654–63. 10.1002/bies.201500192.

Zhu, Lan, and Carlos D. Bustamante. 2005. “A Composite-Likelihood Approach for Detecting Directional Selection From DNA Sequence Data.” Genetics 170 (3): 1411–21. 10.1534/genetics.104.035097.

