## Supplementary Figures & Tables for "Pervasive cryptic selection in the human noncoding genome"

### Supplementary Figure 1

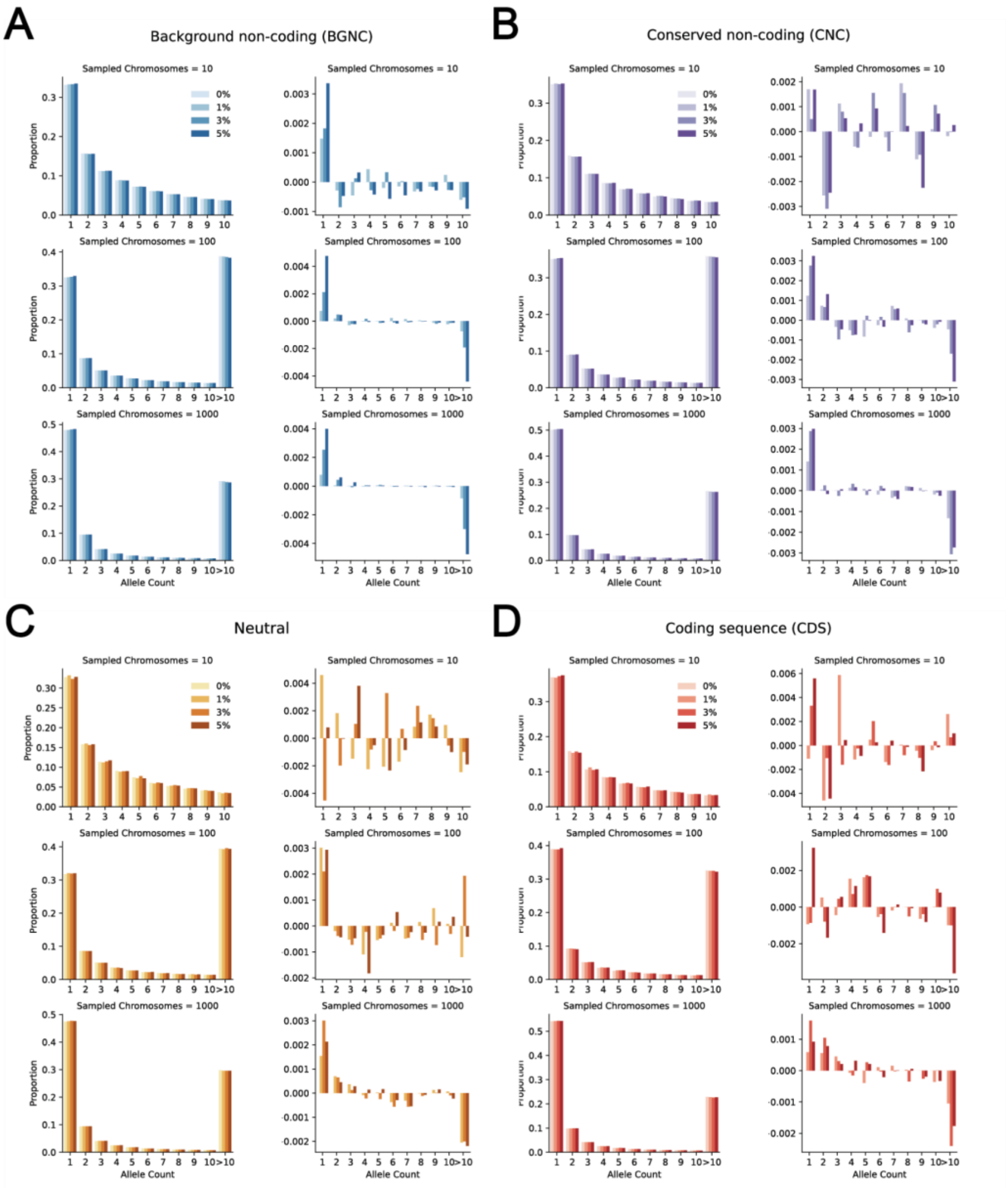


Supplementary Figure 1. SFSs for simple simulations for all region types. In each panel, the first column shows the proportion of variants in each allele frequency bin, and the second column shows the residual by subtracting bin values of 0% from 1%, 3%, and 5%. Rows in each panel correspond to different numbers of sampled haploid genomes. SFSs for BGNC (A), CNC (B), neutral (C), and coding (D) regions are shown here.

### Supplementary Figure 2

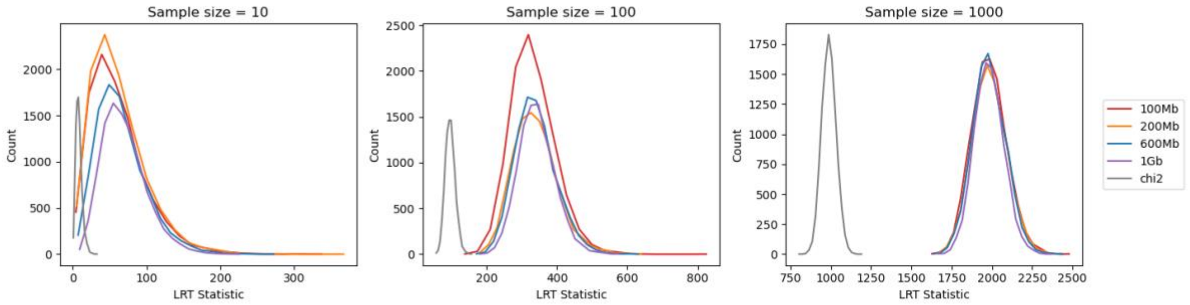


Supplementary Figure 2. Empirical null distributions and corresponding chi-square distributions for simple simulations. The LRT behaves asymptotically chi-square, but the empirical null distributions created by comparing control SFSs to each other are considerably shifted to the right across all three sample sizes. This is likely reflecting the high degree of linkage when assessing several Mb of sequence.

### Supplementary Figure 3

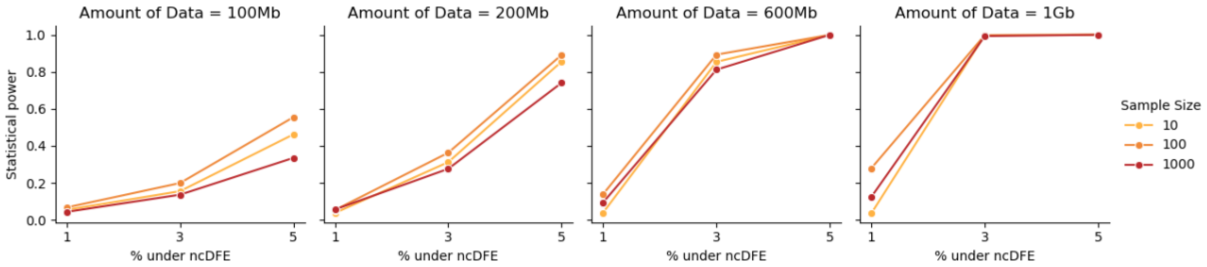


Supplementary Figure 3. Statistical power for LRT analysis of simple simulations from Figure 3. Each panel corresponds to a different amount of sequence data, with lines representing each sample size. Each line consists of three points corresponding to the power at each proportion of mutations under the ncDFE.

### Supplementary Figure 4

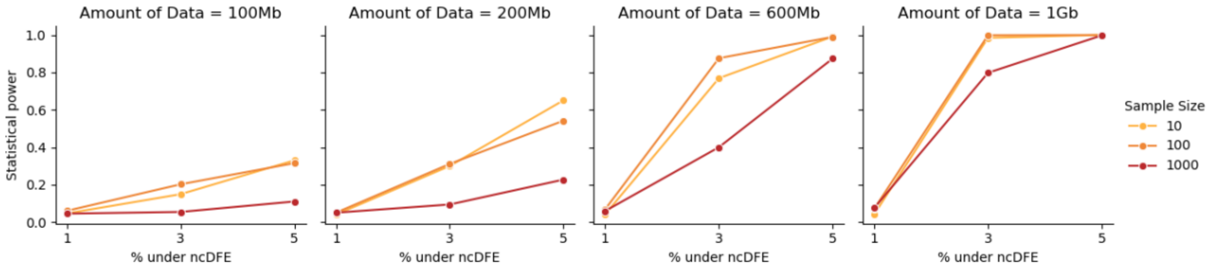


Supplementary Figure 4. Statistical power for LRT analysis of complex simulations from Figure 4. Each panel corresponds to a different amount of sequence data, with lines representing each sample size. Each line consists of three points corresponding to the power at each proportion of mutations under the ncDFE.

Supplementary Figure 5

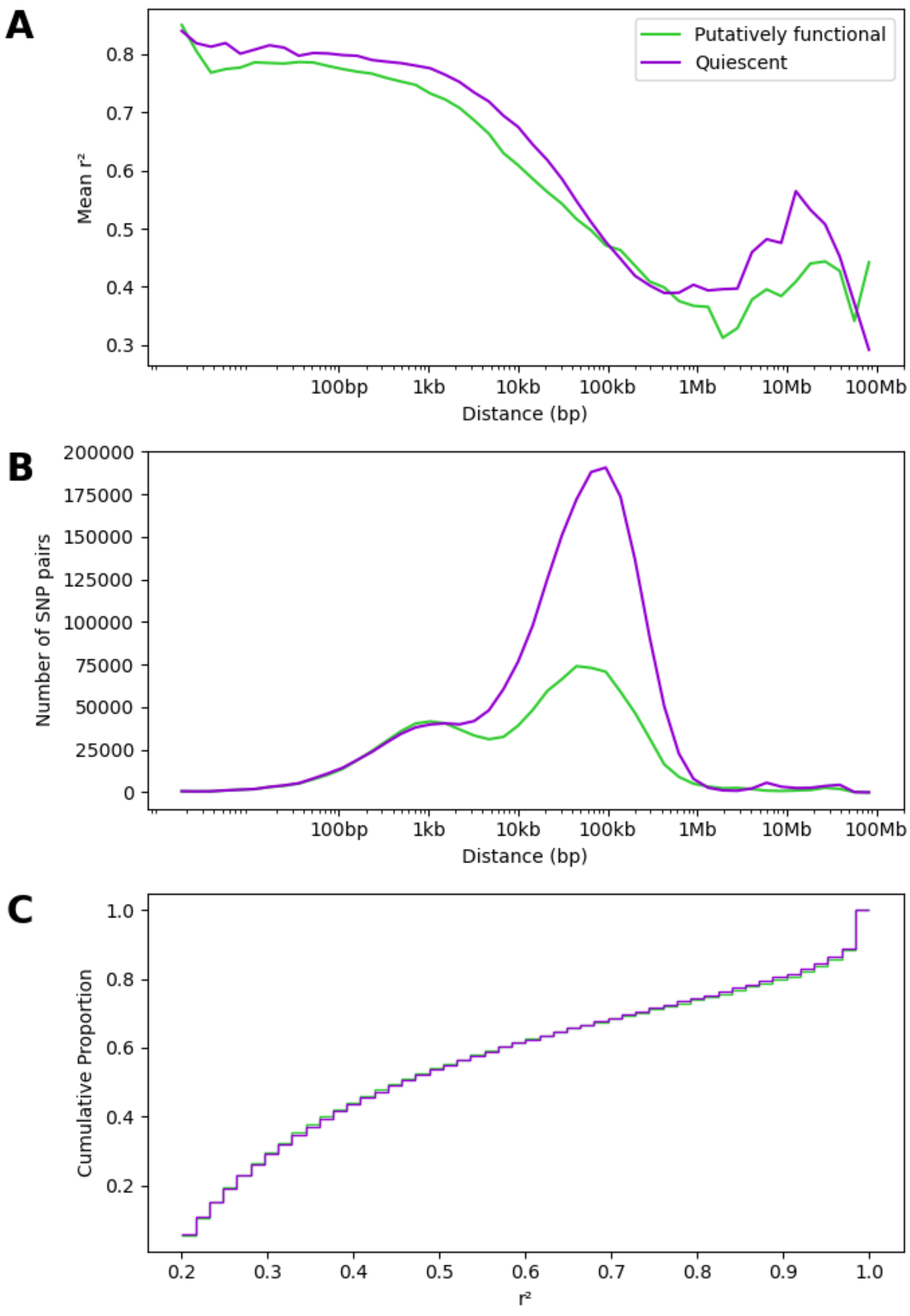


Supplementary Figure 5. Linkage disequilibrium (LD) patterns in 100 Mb subsets of putatively functional and quiescent regions. (A) Mean $r^2$ binned by pairwise SNP distance. (B) SNP pair counts binned by pairwise SNP distance. (C) Cumulative distributions of $r^2$ values for SNP pairs with pairwise distance between 1 kb and 1 Mb.

### Supplementary Figure 6

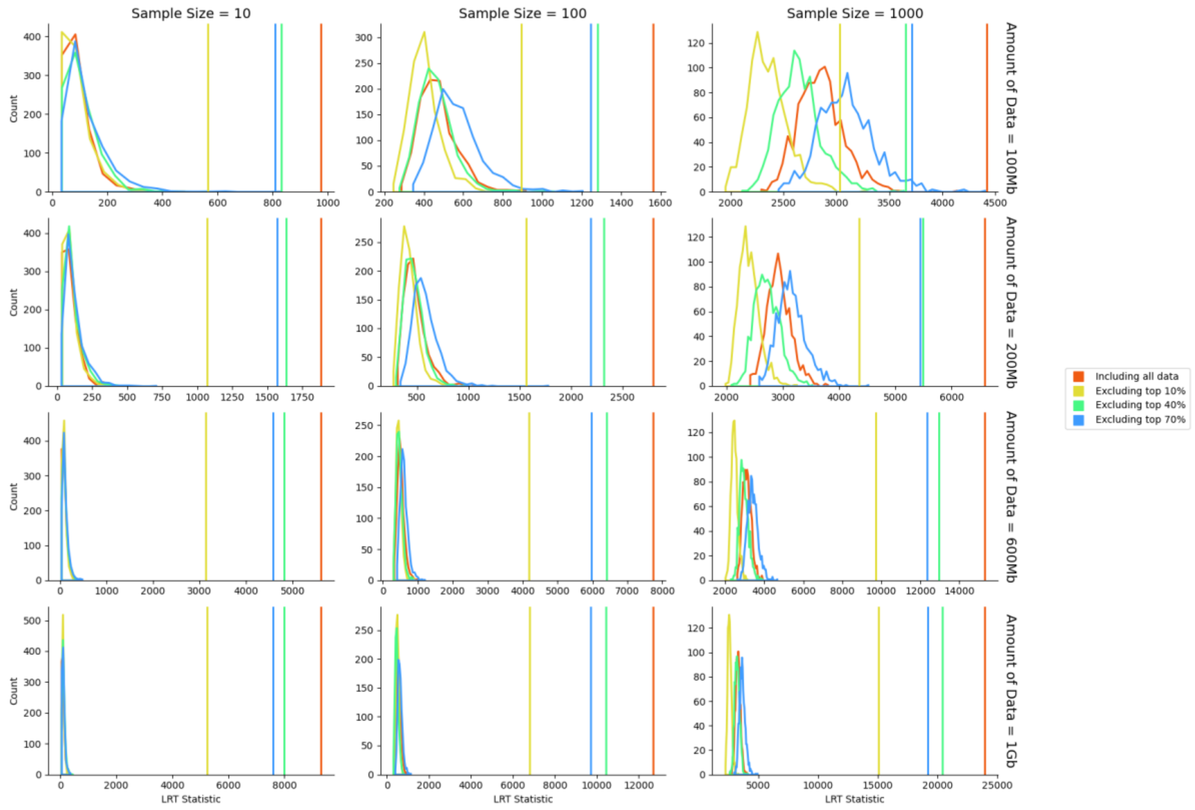


Supplementary Figure 6. LRT distributions for 1KGP data across sample sizes and amounts of tested sequence from Figure 6. Each color corresponds to a different PhastCons filtering threshold. Colored distributions are the null distributions from comparing quiescent SFSs to each other. Solid colored lines denote statistically significant comparisons between putatively functional and quiescent. All comparisons shown here are statistically significant.

### Supplementary Figure 7

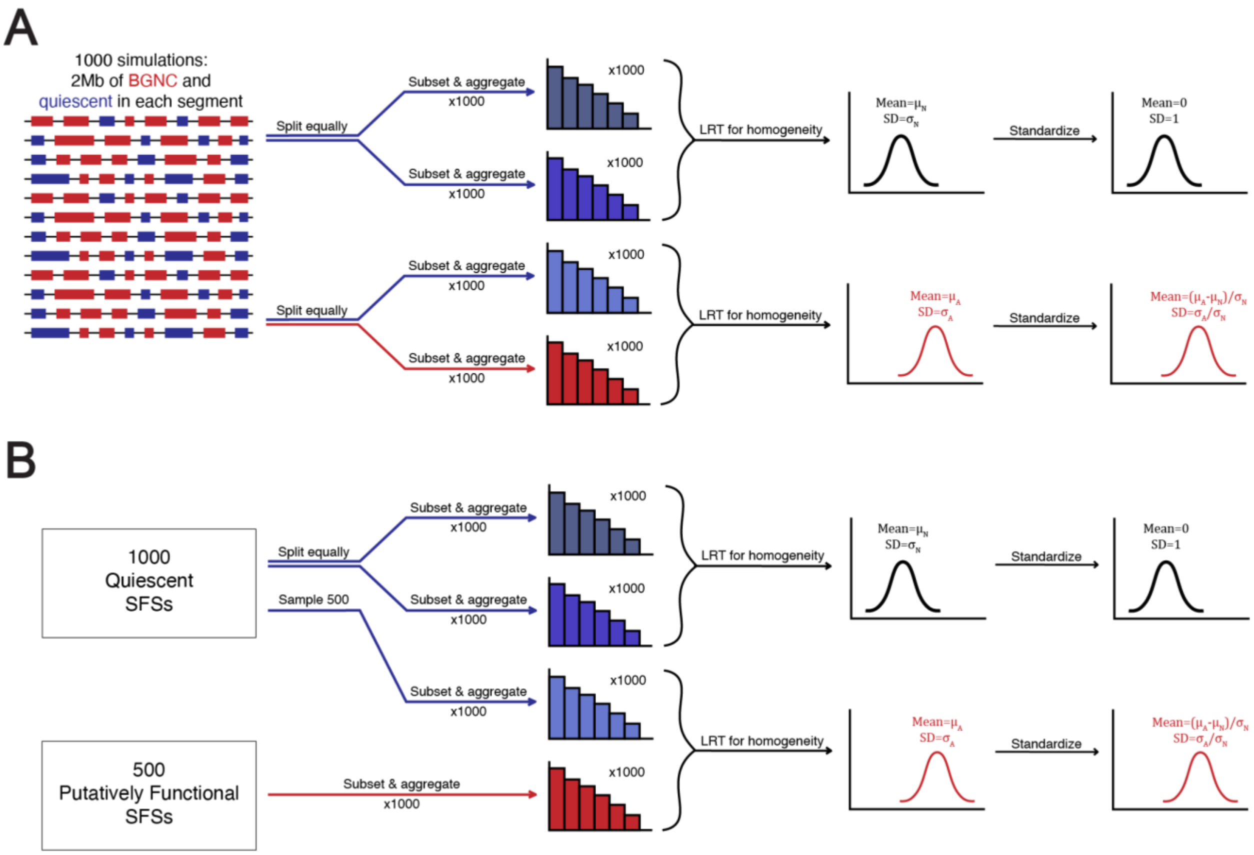


Supplementary Figure 7. Diagram illustrating SFS comparison strategy for SFSs from 1KGP data and matched simulations from Figure 7. (A) Quiescent SFSs extracted from simulations are divided into two equally-sized sets. SFSs within each set are randomly subsampled and aggregated iteratively, and the resulting aggregate SFSs are compared using the LRT of homogeneity to generate the empirical null distribution. The alternative distribution is constructed identically, except quiescent SFSs from one set are compared against BGNC SFSs from the other. Alternative distributions are then standardized using the mean and standard deviation of the corresponding null distribution. (B) Sets of quiescent and putatively functional SFSs are constructed by randomly sampling 2 Mb of each region type from the 1KGP data. To generate the null distribution, quiescent SFSs are divided into two equally-sized subsets and subjected to the same bootstrapping and aggregation procedure, with aggregate SFSs compared using the LRT of homogeneity. The alternative distribution is generated by comparing quiescent and putatively functional SFSs, and is standardized using the mean and standard deviation of the null distribution.

#### Supplementary Table 1

| % BGNC mutations under ncDFE | % BGNC mutations under selection |
|---|---|
| 0 | 0 |
| 1 | 0.261 |
| 3 | 0.782 |
| 5 | 1.30 |
| 10 | 2.61 |
| 20 | 5.21 |
| 40 | 10.4 |
| 60 | 15.6 |
| 80 | 20.86 |

Supplementary Table 1. Percent of mutations under the ncDFE in BGNC regions in simulations converted to the percent of mutations under negative selection. This is calculated by multiplying the left column by 0.26, which is the proportion of mutations with a selection coefficient of at least $10^{-5}$ in the ncDFE (see Methods).

#### Supplementary Table 2

| Top PhastCons sites excluded | Putatively Functional (Mb) | Quiescent (Mb) |
|---|---|---|
| 0% | 1293.45 | 688.32 |
| 10% | 1058.7 | 588.03 |
| 40% | 560.98 | 278.98 |
| 70% | 316.51 | 147.74 |

Supplementary Table 2. The amounts of putatively functional and quiescent sequence after excluding sites within the top X% of PhastCons primate-wide and mammalian-wide scores.

### Supplementary Table 3

| | 0% under ncDFE | 10% under ncDFE | 20% under ncDFE | 40% under ncDFE | 60% under ncDFE | 80% under ncDFE |
|---|---|---|---|---|---|---|
| **10% / 10%** | 5.65% | 7.02% | 8.39% | 11.1% | 13.9% | 16.6% |
| **20% / 20%** | 9.33% | 10.5% | 11.6% | 13.9% | 16.2% | 18.5% |
| **10% / none** | 2.62% | 4.29% | 5.96% | 9.31% | 12.7% | 16.0% |
| **20% / none** | 4.02% | 5.70% | 7.37% | 10.7% | 14.1% | 17.4% |

Supplementary Table 3. The amounts of estimated genome-wide constraint for each simulation data threshold (columns) and each 1KGP data comparison (rows).
